# Interhemispheric transfer of sensory and working memory information is dictated by behavioral strategy

**DOI:** 10.64898/2026.03.18.712808

**Authors:** Elad Avidan, Shlomo David Sherer, Ariel Gilad

**Affiliations:** Department of Medical Neurobiology, Faculty of Medicine, The Institute for Medical Research Israel-Canada (IMRIC), The Hebrew University of Jerusalem, Jerusalem, Israel

## Abstract

The cerebral hemispheres process sensory information largely independently, yet coherent perception and behavior requires rapid interhemispheric communication. Which cortical area transfer information across hemispheres and what governs transfer location remains poorly understood. Here, we combined bilateral whisker-based texture-matching tasks with cortex-wide calcium imaging in mice to investigate the interhemispheric transfer of sensory and working memory information. Mice compared textures presented either simultaneously to both whisker pads or sequentially with a delay, thereby requiring the transfer of sensory or working memory content across hemispheres. Despite identical task demands, animals adopted distinct active or passive behavioral strategies that dictate the cortical location of interhemispheric transfer. Passive mice route both sensory and working memory information through the posterior lateral association cortex, whereas active mice rely primarily on bilateral barrel cortices. During delayed comparisons, active mice rapidly shift information to the contralateral barrel cortex, exhibit impatience, and display reduced performance, indicating that barrel cortices are suboptimal for transferring working memory. These findings identify behavioral strategy as a key determinant of interhemispheric communication and demonstrate a flexible, state-dependent routing of information across the cortex.

## Introduction

Despite the predominantly contralateral processing of sensory inputs, the brain successfully integrates information from both hemifields into a single, unified percept. Achieving such coherence relies on rapid and flexible communication between the two cerebral hemispheres, particularly across cortical areas, and this interhemispheric exchange is essential during natural behavior^1–5^. For example, a mouse navigating a narrow tunnel relies on bilateral whisker input to maintain a continuous representation of the surrounding space, necessitating rapid transfer of sensory information between the hemispheres. Beyond immediate sensory processing, interhemispheric communication is also critical for higher-order functions such as working memory (WM), which involves temporarily maintaining information available for processing^6–8^. A mouse that detects a predator in one visual hemifield and anticipates its reappearance in the other must maintain and transfer information across hemispheres to guide behavior^6,9,10^. Despite its fundamental importance, the principles governing interhemispheric transfer of behaviorally relevant information remain poorly understood^2,4,11^. Specifically, it remains unclear which cortical areas mediate this transfer, which behavioral states permit interhemispheric exchange, and whether different forms of information, sensory versus WM, engage distinct interhemispheric pathways.

The two hemispheres are primarily connected by the corpus callosum (CC), with many cortical areas projecting densely to their homotopic counterparts regions^12–15^. Consistently, resting-state and anesthetized recordings have revealed strong interhemispheric functional connectivity during spontaneous activity^16–19^. In addition, unilateral sensory manipulations induce plastic changes in homotopic cortical regions, suggesting that callosal projections may influence sensory processing^20–31^. However, most of these studies do not explicitly require the transfer or comparison of task-relevant information across hemispheres. Among the few that do, Park et al. (2025) trained mice to compare whisker stimulus locations and observed enhanced interhemispheric coordination between neurons in the bilateral barrel cortices that emerged with learning. Similarly, Brincat et al. (2021) showed that during a WM-guided saccade task in monkeys, working memory traces transfer from one lateral prefrontal cortex to the other when the remembered location crosses hemifields. These findings indicate that both sensory and higher-order information can be transferred across hemispheres in a task-dependent manner, yet a systematic understanding of goal-directed interhemispheric interactions across the cortex is still lacking.

An additional, largely unexplored factor shaping cortex-wide dynamics is the behavioral strategy adopted by the animal. During head-fixed tasks, mice can engage stimuli actively or adopt a more passive sensing strategy^32–34^. We previously showed that in a delayed-response texture discrimination task, short-term memory is maintained in distinct cortical regions depending on strategy: active mice predominantly recruit secondary motor cortex (M2), whereas passive mice maintain memory in a posterior lateral cortical area (P)^32,33^. Memory-related activity in M2 is closely linked to upcoming actions, whereas activity in area P more strongly reflects past sensory identity^32^. Consistent with this, area P has been implicated in multiple higher-order sensory processes, positioning it as a potential hub for sensory information integration.

Based on these observations, we hypothesized that the cortical routes mediating interhemispheric transfer depend on both the nature of the information and the animal’s behavioral strategy. Specifically, we predicted that when WM must be maintained without advance knowledge of the required action, as in a delayed match-to-sample task^35–37^, area P would support WM maintenance and interhemispheric transfer. Due to the strong callosal connectivity between homotopic P regions^38,39^, this area is well suited to facilitate such exchange. In contrast, when rapid transfer of immediate sensory information is required, lower-order sensory areas such as the barrel cortex (BC) may play a dominant role.

To test these hypotheses, we trained mice to perform texture-matching tasks in which stimuli were presented either simultaneously to both whisker pads or sequentially with a delay period, while monitoring dual hemisphere cortical calcium dynamics using wide-field imaging. We found that the cortical pathways supporting interhemispheric transfer were not fixed but instead depended on the behavioral strategy adopted by the mouse. In passive mice, task-relevant information was transferred primarily via area P, whereas in active mice, transfer was mediated by BC. These results identify behavioral strategy as a key determinant of interhemispheric communication and demonstrate that the cortex flexibly routes information across hemispheres according to task demands and internal state.

## Results

To study the interhemispheric transfer of information, we trained mice on two whisker-dependent go/no-go matching tasks that required comparing textures presented to opposite sides (Fig. 1). In the first task (‘BOTH’), the two textures were presented simultaneously, requiring mice to transfer sensory information across cortical hemispheres. In the second task (‘DELAY’), a several-second delay interval separated the two stimuli, requiring the transfer of WM correlates across hemispheres. During task performance, wide-field calcium imaging (GCaMP6f) was employed to monitor neural dynamics across the superficial layers of both dorsal hemispheres (Fig. 2). In total, five male mice were trained on both tasks. Training required 24.6 ± 5.27 sessions for the BOTH task and an additional 15.4 ± 2.94 sessions for the DELAY task (mean ± S.E.M.). The total experimental protocol spanned an average of 184.8 ± 6.1 days per mouse (gross).

**Figure 1.**
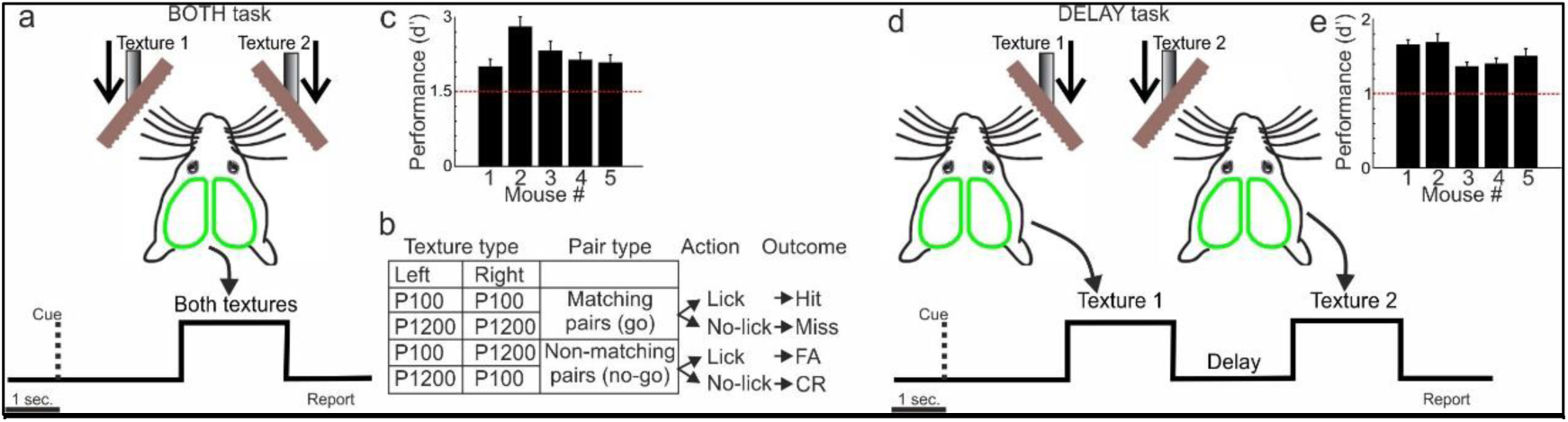
Behavioral tasks and performance. **a.** Schematic illustration of the BOTH task. **b.** Trial types and outcomes. **c.** Performance (d’) in the BOTH task for each mouse. Dashed red line indicate expert threshold. Error bars indicate SEM over recording sessions (n=13, 16, 13, 12 and 11 for mouse 1-5 respectively). **d.** Schematic illustration for the DELAY task. **e.** Performance (d’) as in c but for the DELAY task (n=24, 44, 32, 24 and 26 for mouse 1-5 respectively).

**Figure 2.**
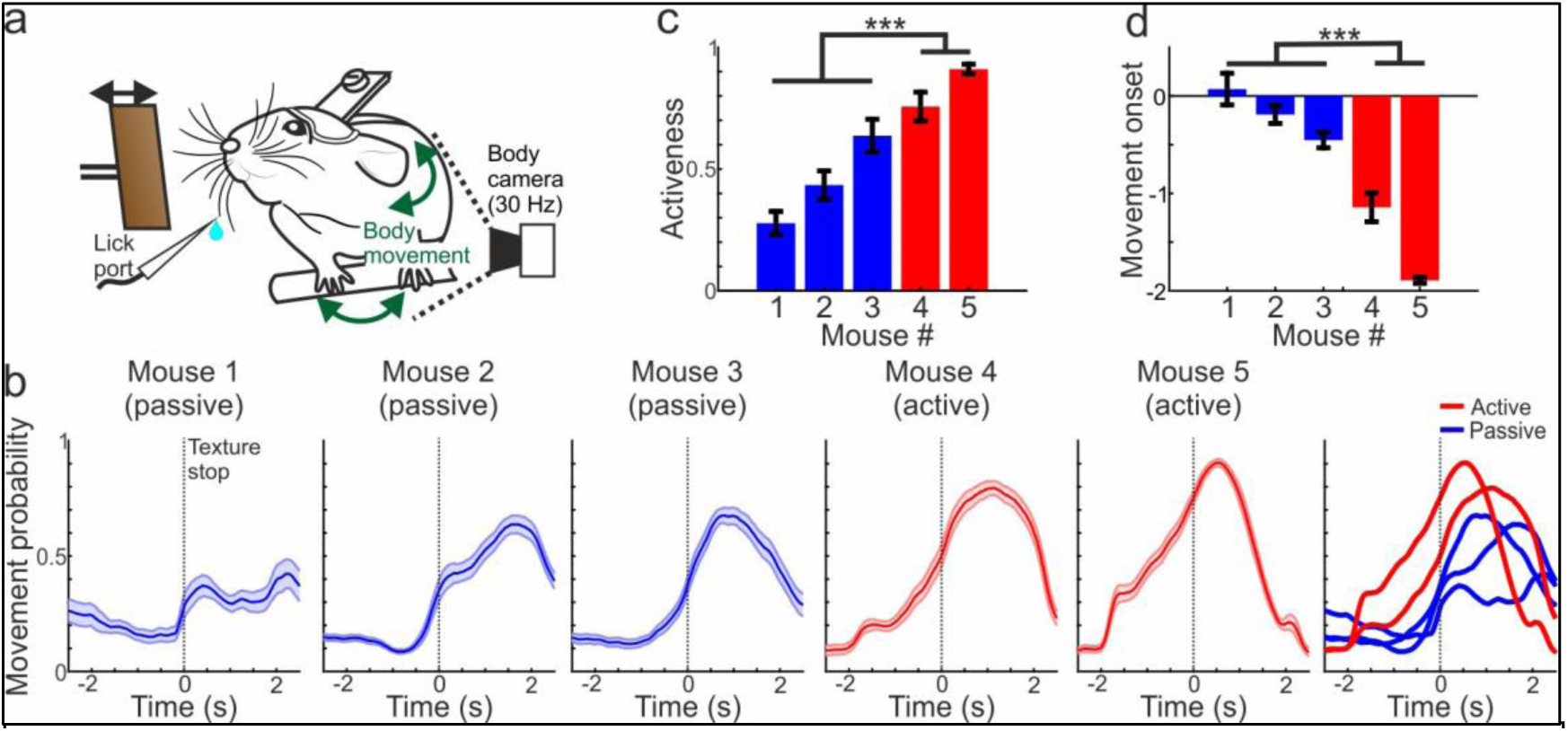
Behavioral strategy in the BOTH task. **a.** Schematic illustration of the behavioral setup and body camera. **b.**Movement probability as a function of time during the BOTH task for Hit trials in each of the five mice. In blue are passive mice and in red are active mice. Error bars depictSEM across recording sessions (n=13, 16, 13, 12 and 11 for mouse 1-5 respectively). On the right are all movement vectors displayed together. **c.**Activeness (i.e., movement probability during the sensation period; -1 to 1 seconds relative to texture stop) for each mouse. Error bars as in b. **d.**Movement onset for each mouse. Error bars as in b. *** -p<0.001; Wilcoxon rank-sum test.

We first focus on BOTH task (Figures 1–4). In short, each trial began with an auditory cue, signaling the simultaneous approach of two sandpaper textures (grit size P100: rough; P1200: smooth) to the whiskers on both sides of the snout (Fig. 1a). The textures presented were either identical (i.e., right: P100 and left: P100; or right: P1200 and left: P1200), or different (e.g., right: P100 and left: P1200; or right: P1200 and left: P100). Matching textures constituted the ‘go’ condition while non-matching pairs served as the ‘no-go’ condition (Fig. 1b). The textures remained in contact with the whiskers for 2 seconds before retraction. Following retraction, mice were rewarded with water for licking during go trials (i.e., Hit trial). In no-go trials, licking responses (false alarms; FA) triggered a mild white noise. No reward or punishment was delivered for correctly withholding licks in no-go trials (Correct Rejections; CR) or for failing to lick in go trials (Miss).

**Figure 3.**
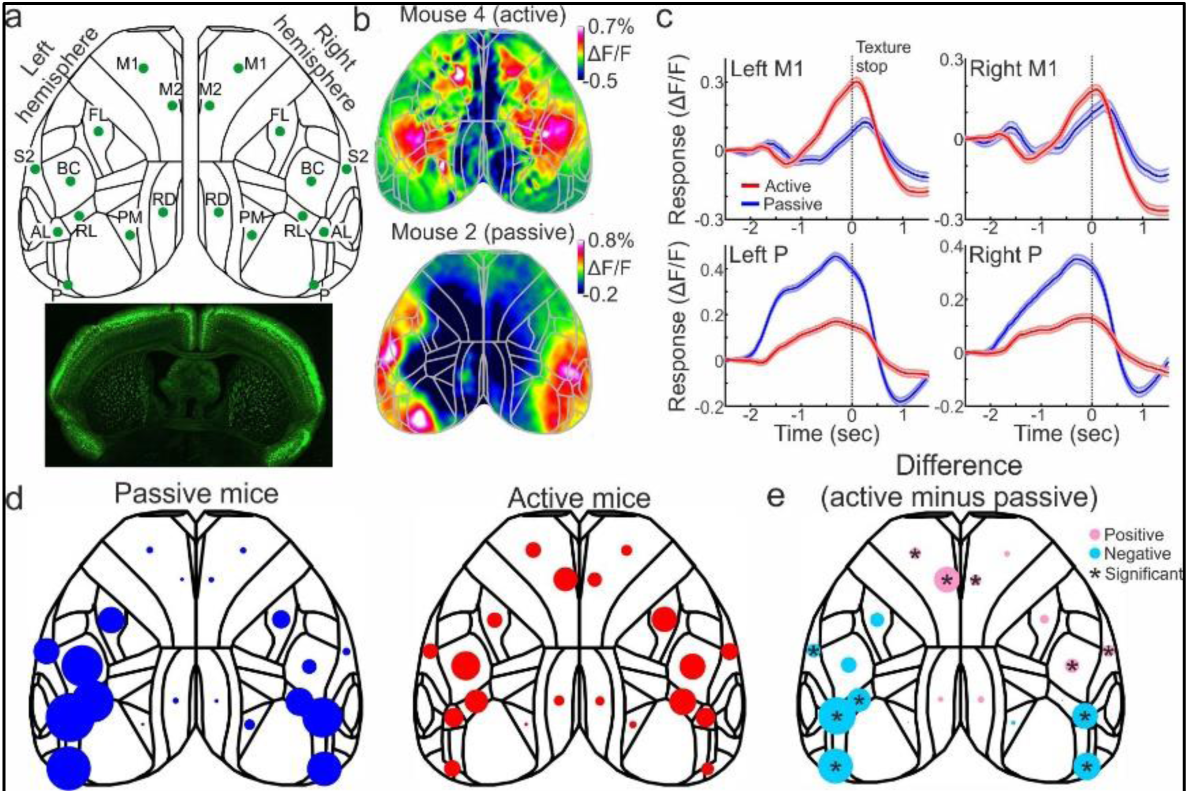
Strategy dependent response profiles in the BOTH task. **a)** Wide-field imaging preparation for imaging the whole dorsal cortex. *Top:* 20 Cortical brain areas were defined, 10 for each hemisphere. P – Posterior, AL – Anterior lateral, PM – Posterior medial, RL – Rostrolateral, RD – Retrosplenial dorsal, BC – Barrel cortex, S2 – Secondary somatosensory, FL – Somatosensory forepaw, M1 – Primary motor, M2 – Secondary motor. *Bottom:* Coronal slice displaying GCaMP6f in superficial layers across the cortex. **b.** Example activity map (color depicts ΔF/F; CR condition) during the sensation period in an active (top) and passive (bottom) mouse. **c.** Temporal responses in four cortical areas (Left M1 and P; Right M1 and P) during the CR conditions in active (red) and passive (blue) mice. Error bars depict SEM across trials from 3 passive mice and 2 active mice (n = 3093 and 1781 trials respectively). **d.** Response profile within the sensation period (-0.5 to 0 relative to texture stop) during the CR conditions for each cortical area in passive (left) and active (right) mice. The bigger the circle, the larger the response (ΔF/F). **e.** Differential response profile across the cortex derived from the two profiles in d (active minus passive). * - p<0.05. Wilcoxon rank sum test. FDR corrected.

**Figure 4.**
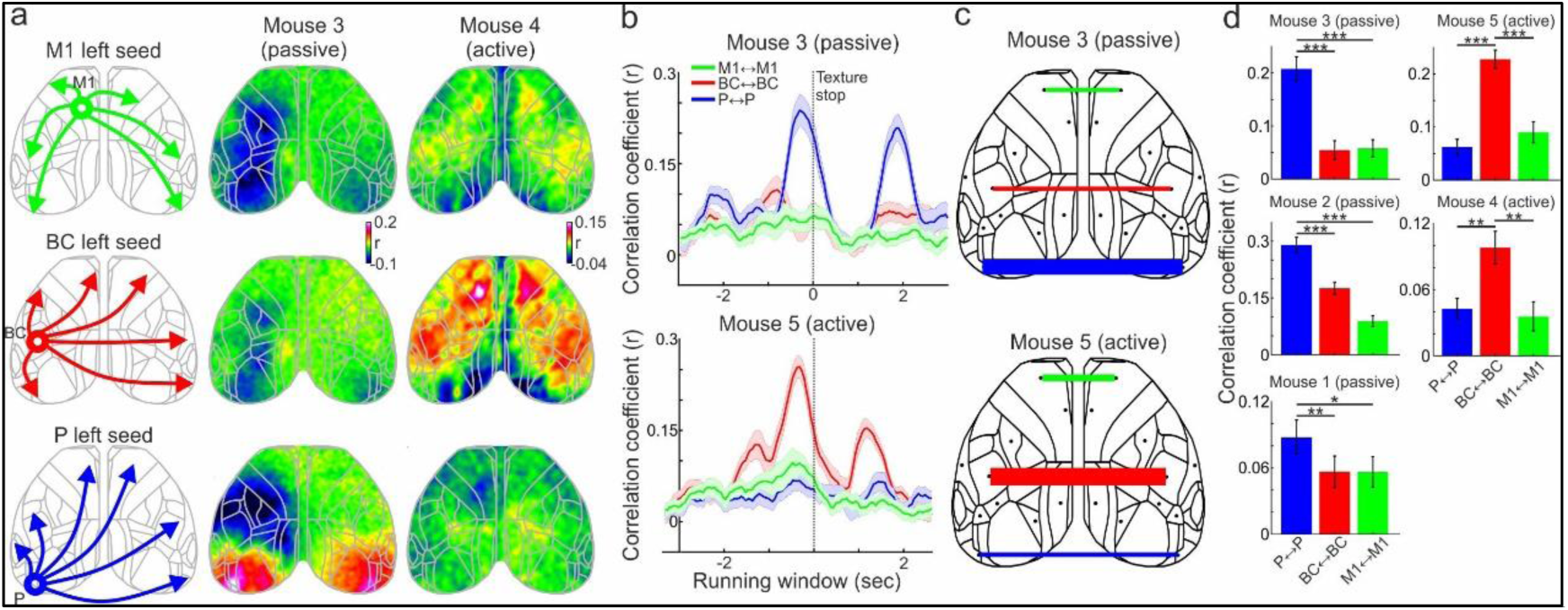
Strategy dependent interhemispheric correlations during the BOTH task. **a.** *Left:* schematic illustration for three different seed correlation maps (for a window during the sensation period) each displaying a different seed area (M1, BC and P). *Middle:* Corresponding seed correlation maps (colors depict r values) in an example session in a passive mouse. *Right:* Example active mouse. **b.** Correlations in a running window (1 sec.) for three interhemispheric pairs (P and P in blue; BC and BC in red; M1 and M1 in green) in an example passive (top) and active (bottom) mouse. Error bars depict SEM across recording sessions (n=13 and 11 for the passive and active mouse). **c.** Correlation values (r) of the three interhemispheric pairs during the sensation period plotted on top of the cortical map for an example passive (top) and passive (bottom) mouse. Thick lines depict high r values. **d.** Correlation values for the three interhemispheric pairs during the sensation period for each mouse. Error bars depict SEM across recording sessions (n=13, 16, 13, 12 and 11 for mouse 1-5 respectively). * - p<0.05. ** - p<0.01. *** - p<0.001. Wilcoxon rank sum test.

Importantly, successful task performance required not only the identification of the texture-type presented to each individual whisker pad but also the integration of sensory information from both hemispheres in order to determine whether the textures matched. Furthermore, the task design enabled a comparison between two distinct ‘Hit’ conditions, P100-P100 vs. P1200-P1200, allowing for the investigation of texture-type recognition. Alternatively, comparing ‘Hit’ vs ‘CR’ trials across different stimulus pairs enabled the study of choice-related encoding. All five mice displayed expert performance in the BOTH task, exceeding a d’ prime of 1.5 (Fig. 1c). We found no substantial performance bias toward any specific stimulus pair (Fig. S1a). An important factor of neuronal dynamics is the behavioral strategy of the mouse, i.e., body movement throughout the trial, which were previously shown to have a profound impact on cortex-wide dynamics^32–34^.

To do this, we calculated the body movement metrics for each trial from the body camera and derived the movement probability as a function of time for each mouse separately (Fig. 2a, b; see Methods). Although the task was identical for all mice, their behavioral strategies (i.e., movement probability vector) differed substantially: three mice were relatively passive, exhibiting late movement onsets (i.e., passive mice 1-3; Video S1), whereas two mice were more active, and with earlier onsets (i.e., active mice 4 and 5; Video S2). To quantify this, we defined ‘activeness’ as the mean movement probability during the sensation period (-1 to 1 relative to texture stop which included the first whisker touches and the early sensation period) and found it was significantly higher in active mice (mice 4 and 5) compared to passive mice (mice 1-3; Fig. 2c; p<0.001; Wilcoxon rank-sum test). In addition, movement onset (i.e., the first frame in which the movement vector exceeded 2STD of the baseline) was significantly earlier in active compared to passive mice (Fig. 2d; p<0.001; Wilcoxon rank-sum test). We further observed that active mice reached expert performance faster in both tasks compared to passive mice (n=37, 31 and 31 sessions for passive mice 1-3; n=11 and 13 sessions for mice 4 and 5), implying that an active strategy is more optimal for the BOTH task, similar to previously reported unilateral tasks^32–34^. A full presentation of the movement profiles for each of the four conditions (two Hit conditions and two CR conditions) is presented in Figure S2. Taken together, although the BOTH task is identical, mice deploy a different behavioral strategy (passive or active) while performing the task.

### Interhemispheric transfer of sensory information is strategy dependent

We next investigated the cortex-wide responses during the BOTH task (Fig. 3a). Activity maps averaged during the sensation period display activity in barrel cortex (BC) with additional frontal responses in the active case and more posterior responses in the passive case (Fig. 3b). Based on these activity maps, we focused on two specific cortical areas: the frontal primary motor cortex (M1) and the posterior association cortex (P). The temporal response during the CR condition in M1 were markedly higher in active mice compared to passive mice in both the left and right hemispheres (Fig. 3c). In contrast, area P display higher responses in passive compared to active mice in both hemispheres (Fig. 3c). To quantify this across the cortex, we defined 10 cortical areas in each hemisphere, i.e., 20 in total, including the somatosensory, association, auditory and motor areas (Fig. 3a). The cortical response during the stimulus period for the CR condition was distributed with a frontal bias for active mice in contrast to a posterior bias for the passive mice (Fig. 3d). The differential response (active minus passive) during CR trials revealed several frontal areas with a significantly positive difference whereas several posterior areas displayed significantly negative differences (Fig. 2b; p<0.001; Wilcoxon rank-sum test; FDR corrected). In addition, we found that most cortical areas displayed significantly higher response in Hit compared to CR trials during the sensation period, due to substantial body movements in Hit trials. A full presentation of the temporal profiles for each of the four pairs and each mouse separately is presented in Figure S3. We note that area P displayed a prominent activity patch in all passive mice substantially before the stimulus period, which was not present in active mice (Fig. S4). Additionally, a control for non-calcium related signals did not show substantial cortical responses (Fig. S5). In summary, beyond the BC, active and passive mice recruit more posterior and frontal cortical areas, respectively.

Next, we investigated the functional relationship between the two hemispheres during the BOTH task. To do this, we first calculated the seed correlation maps during the sensation period for active and passive mice. We selected three cortical areas in the left hemisphere that displayed prominent activity patches: M1 (frontal area), BC (central area) and P (posterior area). In short, we correlated the response during the sensation period (-1 to 1 seconds relative to texture stop) in the seed area with every pixel in the imaging area (Fig. 4a). Each pixel in the map depicts the correlation coefficient (*r* value) with the chosen seed area. Since cortex-wide correlations at the single trial level are often dominated by widespread co-fluctuations, we calculated trial-shuffled correlations where each trial from the seed area was correlated with a random trial from the rest of the cortex (see Methods). These trial-shuffled correlations emphasize temporal dynamics that are time-locked to the task while suppressing non-locked co-fluctuations. Furthermore, we focused this analysis on CR trials, where body movements were substantially smaller during the sensation period (Fig. S2).

In an example passive mouse, the seed correlation map of P (left) displayed high *r* values with the homotopic P area in the right hemisphere (Fig. 4a). In contrast, BC and M1 seed maps exhibited near-zero *r* values with the right hemisphere. Conversely, in an example active mouse, BC seed map displayed higher *r* values with the right BC area and additional frontal areas, whereas P and M1 did not (Fig. 4a). Next, we calculated the correlation between three pairs of homotopic brain areas (P to P, M1 to M1, BC to BC) using a sliding window (1 s) analysis. P to P correlations were high specifically around the sensation period in a passive mouse and BC to BC correlation emerged in the passive mouse (Fig. 4b). This is further displayed in Figure 3c and statistically validated for each mouse separately in Figure 3d. A full presentation of all mice is presented in Figure S6. The full pairwise correlation matrices during the sensation period are presented in Figure S6e. Taken together, we find that interhemispheric correlations are strategy dependent, i.e., via P or BC for passive and active cases, respectively.

Next, we investigated whether cortical dynamics encode texture-type, which is behaviorally relevant. Since the mouse can receive two types of textures to each side (i.e., four different options), there is a crucial need to also discriminate between the type of texture presented. Thus, we hypothesize that key cortical areas such as P and BC, will also encode the type of textures. To do this, we compare between the two Hit trials (P100P100 vs P1200P1200) which were relatively similar in terms of performance and body movements (Fig. S1, 2). First, we present differential activity maps between the two Hit trials within the sensation period (Fig. 5a; P100P100 minus P1200P1200 or vice versa, depending on the mouse). Passive mice displayed prominent differences in one of the P areas, either left or right. In contrast, active mice displayed large differences in both BC areas. We note that in some mice the P100P100 Hit trial was higher whereas in other mice the P1200P1200 Hit trial was higher. We further note that in some cases differential activity shifted in time, for example from left P to right P in a passive mouse.

**Figure 5.**
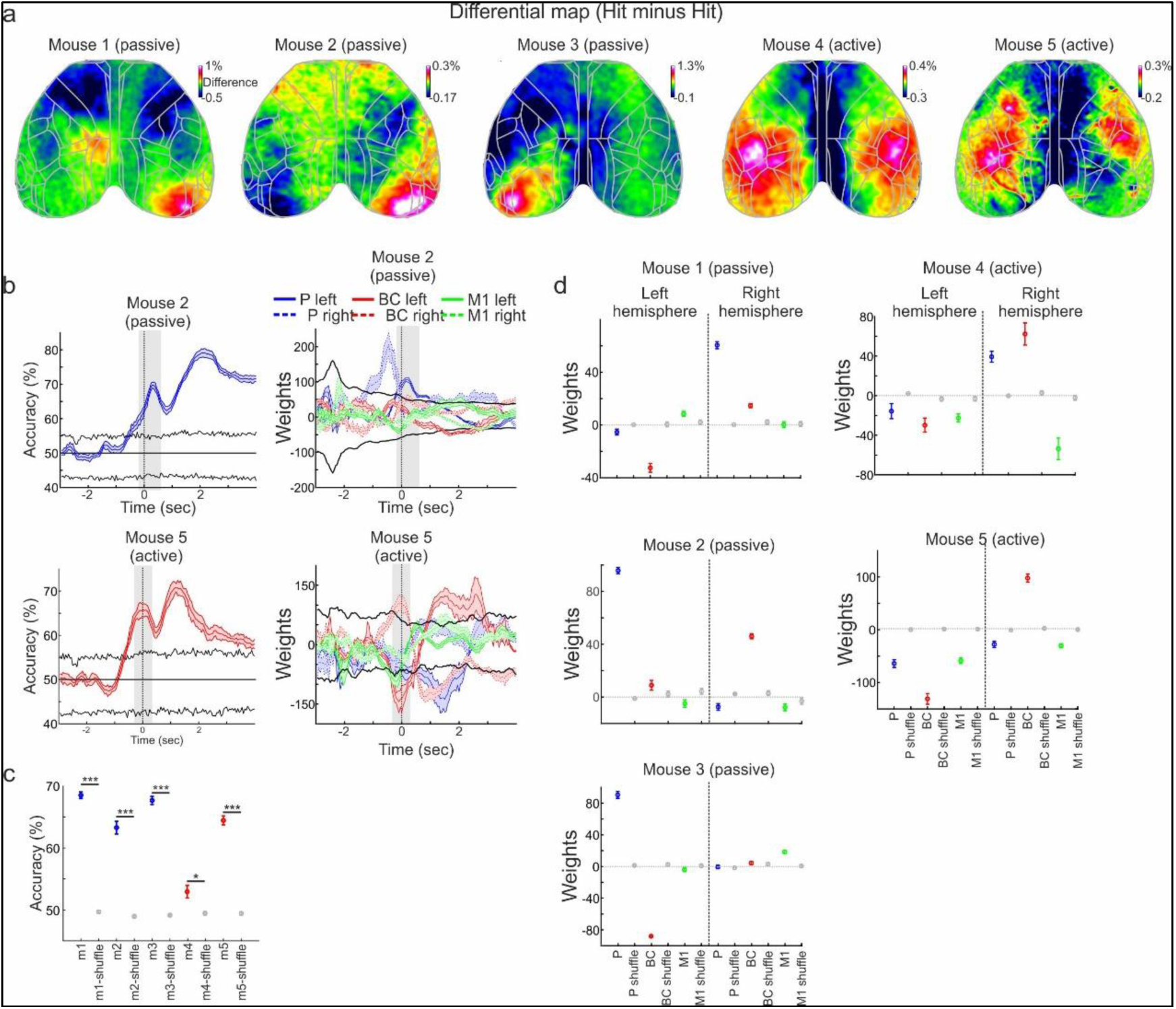
Decoding of texture-type is also strategy dependent. **a.** Example differential response maps for texture-type (subtraction of the two Hit trials where the order is dependent for each mouse) within the sensation period (from -1 to 1 seconds relative to texture stop) for each of the 5 mice. **b.** *Left:* Accuracy as a function of time of an SVM classifier for texture-types (i.e., between two types of Hit trials: P100P100 vs. P1200P1200) for a passive (top) and active (bottom) example mouse. Error bars depict SEM of cross validations (n = 10). Black lines depict mean ± 2*STD of trial shuffled distribution. *Right:* the corresponding weights the classifier assigned to areas P (blue), BC (red) and M1 (green). Solid lines are left hemisphere, and dashed lines are right hemisphere. **c.** Mean accuracy during the sensation period for each mouse. In gray are trial shuffled distributions (n = 100 iterations). Error bars as in b. **d.** Corresponding weights for each mouse and the 6 different cortical areas along with their trial shuffled weights in gray. Error bars as in c. * - p<0.05. ** - p<0.01. *** - p<0.001. Wilcoxon rank sum test.

To investigate the decoding capabilities for the type of texture, we trained a linear Support Vector Machine (SVM) classifier to discriminate between the two Hit conditions based on trial-wise neuronal responses across 20 brain areas (see Methods; 80% train; 20% test; 10 cross validation; Ridge regularization). This was performed separately for each time frame and mouse. The accuracy of the classifier as a function of time exceeded significant levels (crossing 2*std of trial shuffled data; see Methods) around the sensation period in an example active and passive mouse (Fig. 5b). All passive and active mice reached significant accuracies during the sensation period as compared to trial shuffled data (Fig. 5c; p<0.05; Wilcoxon rank-sum test). Nevertheless, the weight distribution assigned by the classifier for active and passive mice was different. The classifier for passive mice assigned the P areas with meaningful weights (i.e., exceeding shuffled boundaries) as compared to M1 and BC (Fig. 5d; left and right). Interestingly, in this passive mouse, the weight shifted from the right P to the left P around the sensation period, implying a sequential transfer of type information. In contrast, in an example active mouse the classifier assigned meaningful weight to both BCs with opposite signs, indicating that one BC favored one texture-type whereas the other favored the other. The weights for the six areas during the sensation period for each mouse emphasizes the difference between passive and active mice, using P and BC respectively (Fig. 5d). In general, during the sensation period, the classifier assigned significant weights to only one of the P areas in passive mice, in two mice the left P and in one mouse the right P. This means that in passive massive the information about the type task was unilateral. In active mice, both BCs were assigned with non-zero and opposite weights, displaying bilateral decoding capabilities. The full details of accuracies and weights for each mouse and all cortical areas are presented in Figure S7. In summary, we find that when simultaneously presented with two textures, passive mice utilize area P to transfer sensory-related information across hemispheres, whereas active mice recruit both BCs instead.

### Working memory transfer across hemispheres is also strategy dependent

Next, we investigated whether WM, the active maintenance of information in the absence of external stimuli, is also transferred across hemispheres via similar pathways. To address this, the same five mice were trained on a DELAY task, in which the two texture presentations were separated by a temporal delay of several seconds (typically 2 seconds; see Methods). Expert performance (*d’*>1 in this task) was slightly but significantly better in passive mice compared to active mice (Fig. 6a; Based on the BOTH task; p<0.05; Wilcoxon rank-sum test). Interestingly, by examining the body movement of each mouse during the DELAY task (for Hit trials), we found that passive mice managed to patiently not move during most of the delay period in anticipation of the incoming stimulus to the other whisker pad (Fig. 6b; Video S3). In contrast, active mice started to vigorously move at the beginning of the delay period and continued to move throughout its duration (Fig. 6c; Video S4). To quantify this, we defined an ‘impatience index’ as the relative movement probability during the early delay period compared to the movement probability during the first texture presentation (see Methods). The impatience index was significantly higher in active compared to passive mice (Fig. 6d; p<0.001; Wilcoxon rank-sum test). Furthermore, active mice exhibited a decline in performance with longer delay periods (>2 seconds), whereas passive mice maintained high performance even with delays of up to 4 seconds. Nevertheless, impatience was found to be significantly and positively correlated with task performance on a session-by-session basis, particularly in active mice (Fig. 6e), indicating that active mice can perform the DELAY task. In summary, active mice became impatient when introduced with a delay period, whereas passive mice remained more still.

**Figure 6.**
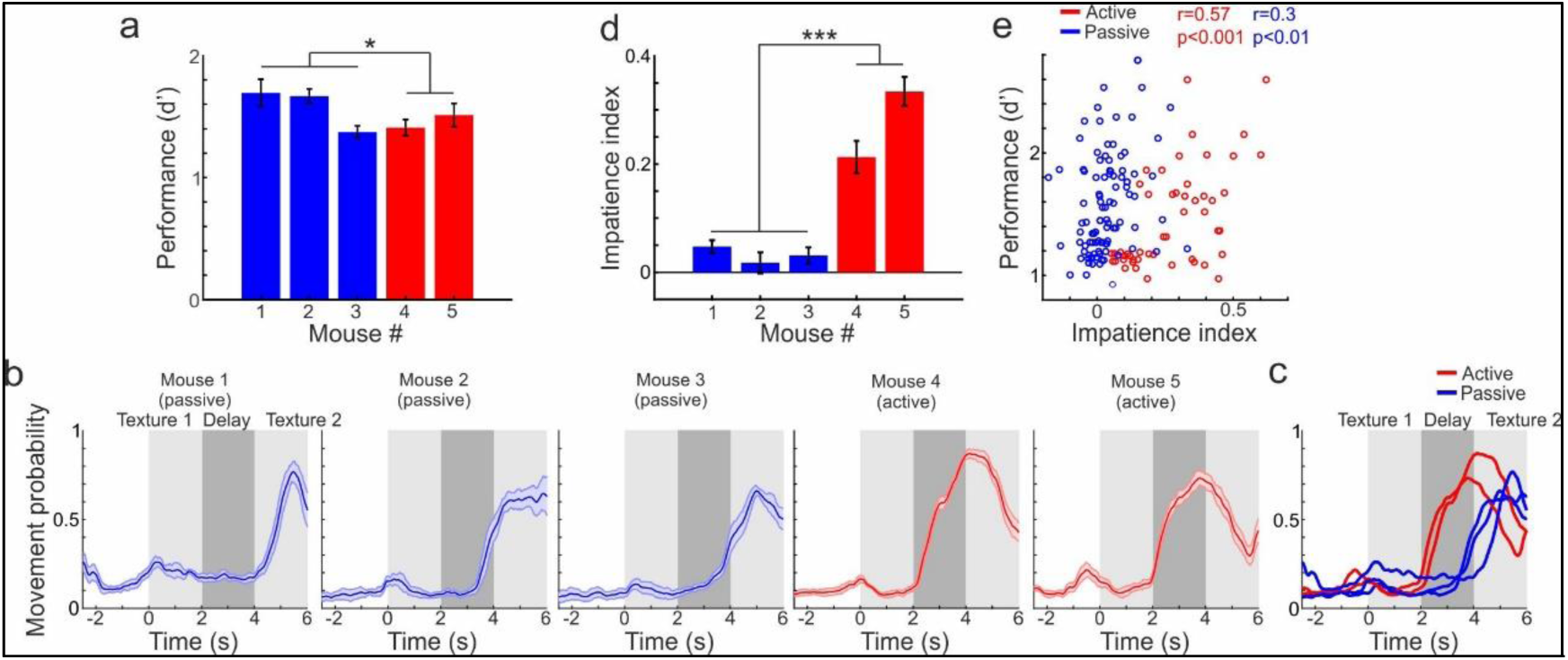
Behavioral strategy in the DELAY task. **a.** Performance (*d’*) in the DELAY task (as in Fig, 1e) but color coded to passive (blue) and active (red) mice. **b.** Movement probability as a function of time during the DELAY task for Hit trials in each of the five mice. In blue are passive mice and in red are active mice. Error bars depict SEM across recording sessions (n=24, 44, 32, 24 and 26 for mouse 1-5 respectively). **c.** All movement vectors displayed together. **d.** Impatience index (i.e., movement probability during the first second of the delay; see Methods) for each mouse. Error bars as in b. **e.** Scatter plot of impatience index (x-axis) versus performance (y-axis). Each dot represents a recording session. Passive and active mice are in blue and red respectively. * - p<0.05. *** - p<0.001. Wilcoxon rank sum test.

Building on these behavioral differences, we next investigated cortex-wide activity maps during the DELAY task. In passive mice we defined four distinct time periods: (1) first texture presentation, (2) early delay period, (3) late delay period and (4) second texture presentation (Fig. 7a). For active mice, since they started to vigorously move during the delay period (resulting in widespread neuronal responses), we focused on three time periods: (1) early sensation of the first texture, (2) later sensation of the first texture and (3) just prior the delay period. In an example session from each passive mouse (CR condition), activity patches started from left BC during the first texture presentation (1), following a shift to an activity patch in the left P area during the early delay period (2) which then shifted to the right P area during the late delay period (3) and finally converged on the right BC during the second texture presentation (4; Fig. 7b top). A video of the activation map as a function of time for a passive recording session highlight the flow of information between hemispheres (Video S5). In contrast, for the two active mice, activity commenced in the left BC during the early sensation period (1) and rapidly shifted to the right BC during the late sensation period of the first stimulus (2; Fig. 6b bottom). Just prior to the start of the delay period (3), activity is maintained in the right BC and, in several instances, propagated to the right P area. The temporal response of these four key areas (left and right P; Left and right BC) revealed that in passive mice, there is a shift from the left P to the right P during the delay period, whereas in the active mouse activity switches from the left BC to the right BC and P areas well within the first texture presentation (Fig. 7c; See also Video S6 for an active mouse example activity maps as a function of time). The activation maps in Figure 3b for active mice during the prior delay period display activity mainly in the right P area, additional examples showing activation patches in both the right P and BC are presented in Figure S8. The grand average across all 20 brain areas in passive mice demonstrated that during the early delay period significant activity was confined to the left posterior association areas (including left P). Subsequently, during the late delay period, activity significantly shifted to the right posterior association areas (including right P; Fig. 7d top; p<0.05 for each presented circle; Wilcoxon sign-rank test; FDR corrected). In active mice, during the early sensation period of the first texture, significant activity emerged in left BC as well as in both somatosensory forelimb cortices (Fig. 7d bottom; p<0.05 for each presented circle; Wilcoxon sign rank test; Bonferroni corrected). Just before the delay period, activity significantly shifted to the right hemisphere including whisker-related sensory areas such as BC, S2 and RL. At this time, responses in the left posterior cortex (including BC and P) were significantly suppressed in active mice. The full temporal profiles for all cortical areas and each mouse are presented in Figure S3b. A control for non-calcium dependent signals is presented in Figure S5c, d. We highlight that in this task, there is no texture presented at the left whisker pad (i.e., corresponding to the right hemisphere), implying that right-hemisphere responses during the first presentation may be driven by the stimulus presented in to the ipsilateral whisker pad.

**Figure 7.**
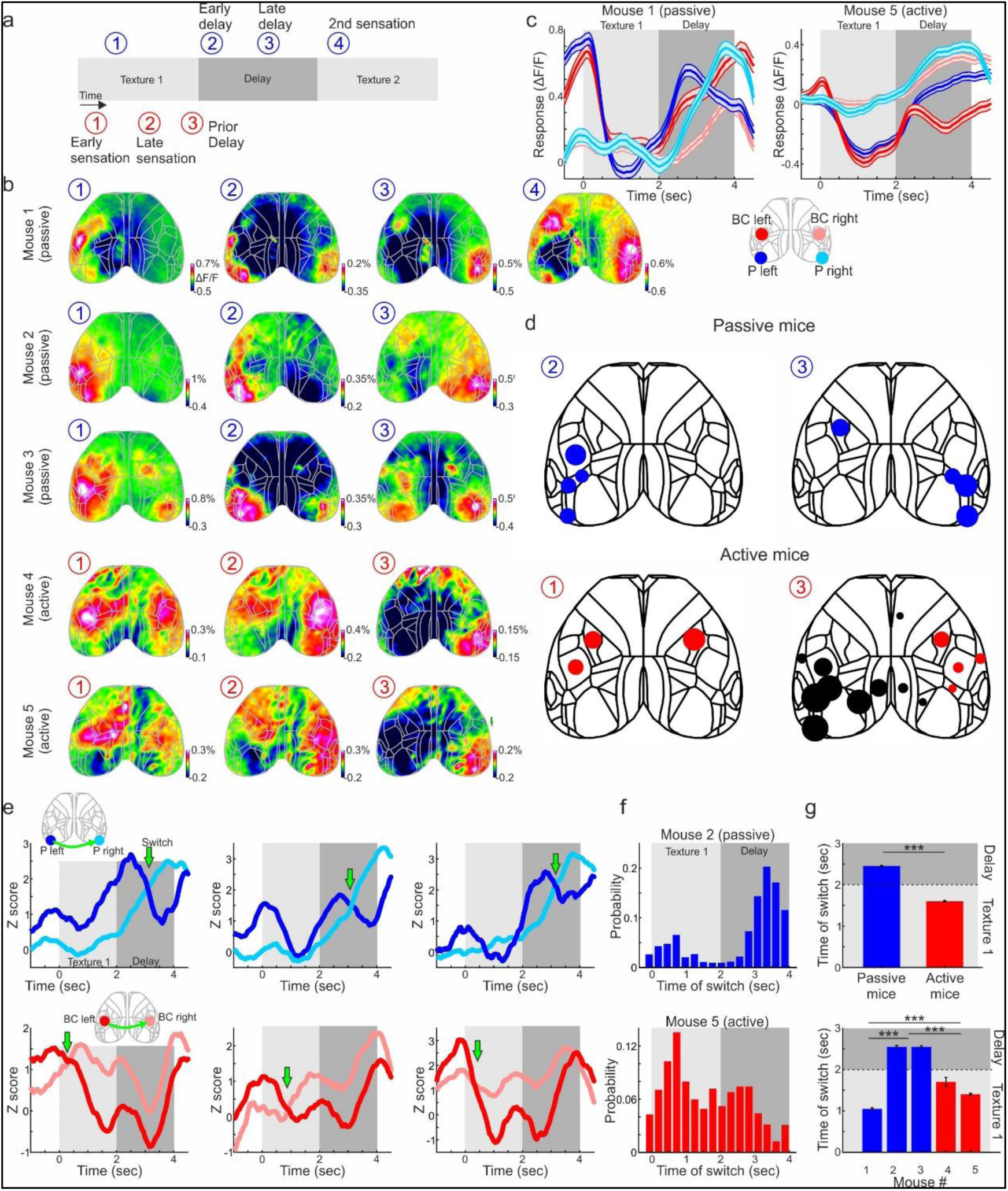
Interhemispheric transfer of working memory during the DELAY task. **a.** Definition of specific time periods during the DELAY task for passive (blue) and active (red) mice. **b.** Activity maps (ΔF/F during CR conditions) during the time periods in *a* for an example recording session for each passive and active mouse. Time point 4 is presented only for mouse 1. **c.** Temporal responses during the CR conditions in four central areas (left and right BC in red shades; Left and right P in blue shades) for an example active and passive task. Error bars depict recording sessions (n= 32 and 26 the passive and active mouse respectively). **d.** Response profile within specific time period (refer to *a*) during the CR conditions for each cortical area in passive (top) and active (bottom) mice. The bigger the circle, the larger the response (ΔF/F). Black circles indicate negative ΔF/F. **e.** Example single trial temporal responses of left and right P (blue shades) for passive mice and left and right BC (red shades) for active mice. Green arrows mark the time of switch between left and right cortical area. **f.** Trial distribution (Hit and CRs pooled together) of the time of switch for an example passive (top) and active (bottom) mouse. **g.** Average time of switch for passive (blue) and active (red) mice pooled together (top) or separately for each mouse (bottom). Error bars are SEM across trials (n=6601 and 3517 trials for passive and active mice respectively; n=2361, 1752, 2488, 1816 and 1701 for mouse 1-5 respectively). * - p<0.05. ** - p<0.01. *** - p<0.001. Wilcoxon rank sum test.

Next, we quantified the time of interhemispheric switch in response during the DELAY task. Single trial examples illustrate the temporal relationships between the bilateral P areas in passive mice and the bilateral BC areas in active mice (Fig. 7e). Intuitively, the switch between the right and left hemisphere might occur at the intersection between the responses of the two areas, as the left area declines after its peak activity and the right area increases response towards its own peak activity (green arrows in Fig. 7e). Based on this, we calculated the moment of activity switch between the two P areas in passive mice and the two BC areas in active mice. In short, the switch was defined as the intersection between the two areas given that both areas had substantial responses within that time (see Methods). This was done for each trial separately. The distribution of switch times in a passive example mouse displays a central peak well within the delay period, whereas the switch distribution in an example active mouse was much earlier, during the first texture presentation (Fig. 7f). Across all mice, the switch time for passive mice was 2.45 ± 1.21 seconds (within the delay period) and 1.6±0.95 seconds for active mice (median ± MAD) with a significant difference between them (Fig. 7g; p<0.001; Wilcoxon rank-sum test). When separating switch times for each mouse, we report that one passive mouse had very early switch times due to a large activity in the right P area early during the first sensation period (Fig. 7g bottom). In summary, passive mice transfer working memory across hemispheres via areas P and during the delay period. In contrast, active mice have trouble in transferring working memory across hemispheres and instead transfer information between BC areas early during the sensation period and start to impatiently move during the delay.

The activity flow described above was consistent across all stimulus types and choices. However, to successfully perform the DELAY task, the mouse must encode the identity of the first texture and maintain it in working memory for subsequent comparison with the second texture. Thus, the ‘type’ information of the first texture must be sustained and presumably transferred to the contralateral hemisphere. To investigate this interhemispheric transfer of information, we trained a machine to classify between the types of texture (Hit P100P100 vs Hit P1200P1200) for each frame separately, similar to the BOTH task (Figure 5). Classifier accuracy exceeded significant levels in both active and passive mice, starting slightly after the first texture presentation and further increasing during the delay period (Fig. 8a). All mice, active and passive, displayed a significant accuracy during the early delay period (first second of the delay) as compared to the accuracy from a trial shuffled data set (Fig. 8b; p<0.001; Wilcoxon rank-sum test). Next, we investigated the assigned weights of the classifier. In an example passive mouse, the classifier first assigned a positive weight to the left P area during the first sensation period, then shifted its weights towards the right P area as the delay started after which the right BC was assigned a significant weight as the second texture approached and touched the whiskers (Fig. 8c). In contrast, in an active mouse example, the classifier assigned significant weight to the left BC which then exhibited a subtle shift to the right BC early during the delay period, but not as prominent as the passive mouse example. Averaged within the early delay period (first second), the classifier assigned significantly non-zero values to bilateral P areas in passive mice and to bilateral BC areas in active mice (Fig. 8d; p<0.05; Wilcoxon rank-sum test as compared to trial shuffle). We note that other areas in certain mice were also assigned with significant weights, for example M1 right in active mouse 5. The full presentation of accuracy and weights of all mice is presented in Figure S9. In summary, information regarding the type of texture flows across hemispheres primarily via P or BC in passive and active mice respectively.

**Figure 8.**
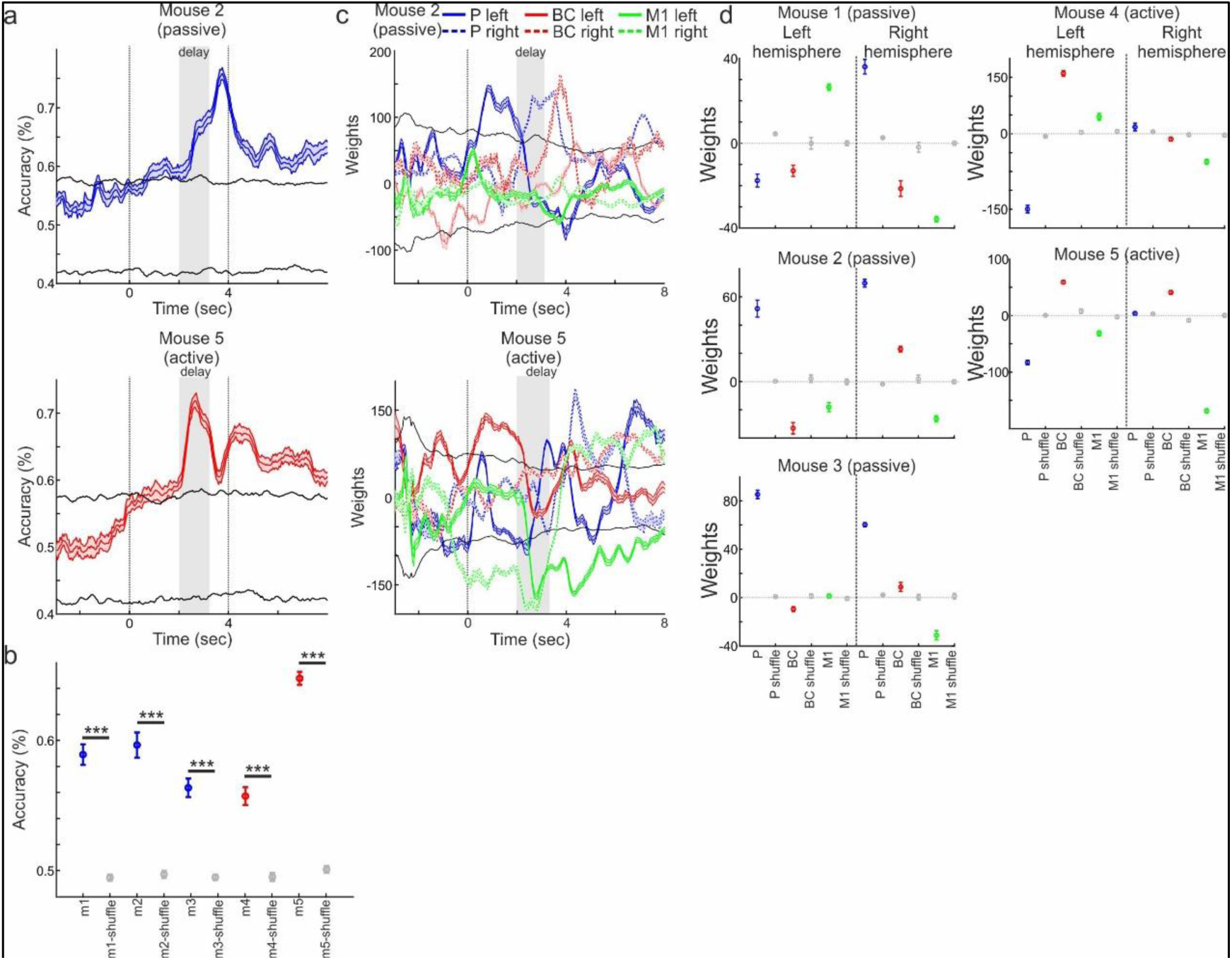
Decoding of texture-type during the DELAY task. **a.** Accuracy as a function of time of an SVM classifier for texture-types (i.e., between the two Hit trials: P100100 vs. P12001200) during the DELAY task for a passive (top) and active (bottom) example mouse. Error bars depict SEM of cross validations (n = 10). Black lines depict mean ± 2*STD of trial shuffled distribution. **b.** Mean accuracy during the early period for each mouse. In gray are trial shuffled distributions (n = 100). **c.** The corresponding weights (from *a*) the classifier assigned to areas P (blue), BC (red) and M1 (green). Solid lines are left hemisphere and dashed lines are left hemisphere. Error bars as in *a*. **d.** Average weights during the early delay period for each mouse and the 6 different cortical areas along with their trial shuffled weights in gray. * - p<0.05. ** - p<0.01. *** - p<0.001. Wilcoxon rank sum test.

## Discussion

In this study, we investigated the interhemispheric transfer of behaviorally relevant sensory information that is either immediately available or must be maintained in working memory (WM). Using bilateral whisker-based texture-matching tasks, we found that sensory information and WM content are transferred across hemispheres via distinct cortical hubs that depend on the behavioral strategy adopted by the mouse. Although task demands were identical, animals differed markedly in their strategies: passive mice transferred information predominantly via posterior lateral association cortex (P), whereas active mice relied mainly on bilateral barrel cortices (BC) for interhemispheric communication (Fig. 9).

**Figure 9.**
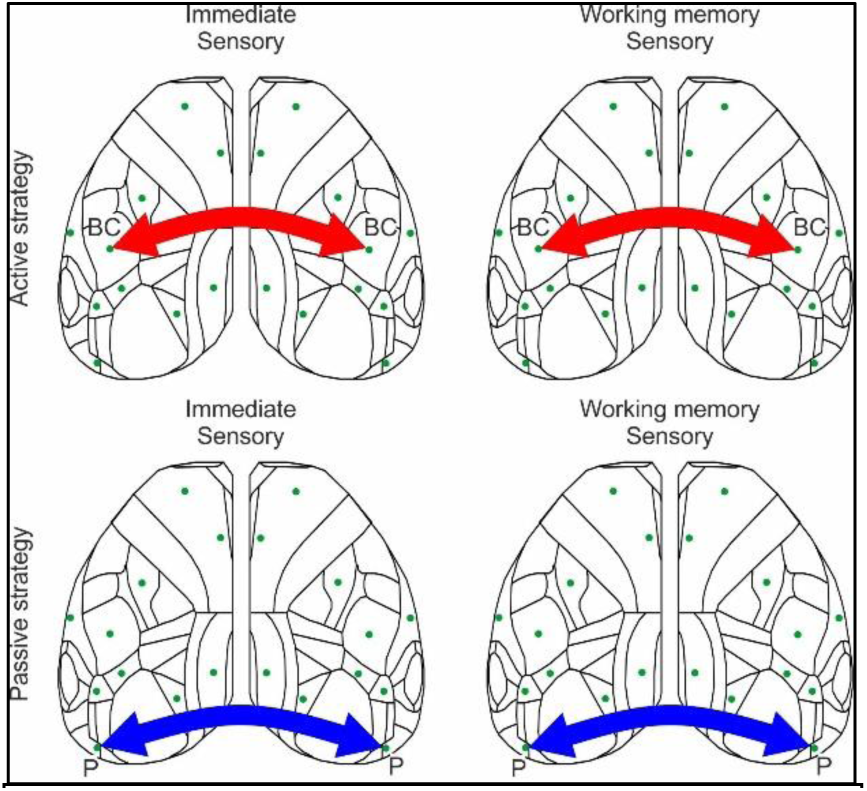
Summary for strategy dependent interhemispheric transfer. Interhemispheric transfer of both Immediate sensory(left)information and WM(right)is via BC for active mice(top;red)and via P in passive mice(bottom;blue).

### Flexible transfer of lower- and higher-order sensory information across hemispheres

Although sensory information is processed primarily in the contralateral cortical hemisphere, effective perception and decision-making require the transfer and integration of information across both hemispheres^2,4,11^. Anatomical and functional studies have identified the corpus callosum, which directly connects the two cortices, as a central substrate for interhemispheric communication^5,17,40,41^. Callosal projections are largely homotopic^12,13^, yet callosal function is highly dynamic and can be modulated by unilateral perturbations, learning, and behavioral demands^3,6,14,22,29–31,42,43^.

Thus, the complexity of the information being transferred appears to dictate the cortical site of interhemispheric exchange. Sensory information requiring immediate transfer may be routed through lower-order sensory areas such as BC, whereas higher-order information, such as WM content, may recruit associative cortical regions. Consistent with this view, callosal neurons are predominantly excitatory and innervate both excitatory and inhibitory targets across multiple cortical layers, supporting flexible and context-dependent interhemispheric communication^44^.

We found that lower-order sensory information was often transferred via BC, however, notably, in passive mice even immediate sensory transfer occurred through the higher-order association area P. For WM, P emerged as an effective hub for interhemispheric transfer, whereas BC was comparatively suboptimal. This observation aligns with previous findings showing that BC is poorly suited for maintaining WM in unilateral tasks^32,33,45,46^. Mice that nevertheless relied on BC for WM transfer tended to do so early, during the first texture presentation, rather than maintaining information across the delay. These animals exhibited lower behavioral performance, initiated impatient and uninstructed movements early in the delay period, and appeared to rely partially on post-transfer P to transiently maintain WM for a limited duration.

Interestingly, the cortical route of interhemispheric transfer, either P or BC, remained stable across task contexts. Mice that transferred information via a particular area during the simultaneous (BOTH) task continued to use the same route during the delayed (DELAY) task, even when that route became suboptimal for WM transfer. This suggests that interhemispheric routing reflects a long-term behavioral strategy rather than flexible, trial-by-trial optimization.

We further found that interhemispheric transfer was localized to a single cortical area, either BC or P, rather than being distributed across multiple regions. This contrasts with intra-hemispheric processing, which can be distributed across networks, and highlights a distinction between binary interhemispheric integration and distributed unilateral computation^7,46–51^. In addition, information about texture identity was encoded bilaterally in BC but unilaterally in P in which representations shifted from one hemisphere to the other. These findings support a division of labor in which BC enables rapid bilateral comparison of immediate sensory input, whereas P mediates sequential, higher-order information transfer across hemispheres.

Together, our findings highlight the flexible nature of interhemispheric information transfer, mediated by the corpus callosum. BC appears well suited for immediate and ongoing sensory transfer, whereas P is optimized for the sequential transfer of higher-order information such as WM. Importantly, this flexibility allows cortical areas to transfer non-canonical information types when dictated by behavioral strategy, even if such routing is suboptimal for task performance.

### Behavioral strategy dictates the cortical location of interhemispheric transfer

Although all mice were trained on identical tasks, they adopted distinct behavioral strategies. During the BOTH task, some mice employed a passive strategy, remaining mostly immobile and allowing the textures to contact their whiskers through passive touch. In contrast, other mice adopted an active strategy characterized by vigorous whisking and body movements initiated shortly after the start cue, resulting in active exploration of both textures.

Previous studies have shown that active strategies can enhance performance and recruit frontal cortical regions, particularly the secondary motor cortex (M2), during unilateral tasks^32–34^. Passive strategies, in contrast, tend to recruit area P, which is engaged in tasks requiring sensory integration within a hemisphere. In the BOTH task, passive mice exhibited strong correlations between neuronal dynamics in the two P areas, whereas active mice showed correlated activity between the two BCs. Thus, even in active mice, the frontal cortex was not used to mediate interhemispheric sensory transfer.

Introducing a delay between the two textures revealed a limitation of the active strategy. Mice that were active during the BOTH task struggled to remain motionless during the delay, exhibited frequent early licks, and showed reduced performance. Increasing the delay duration beyond two seconds proved difficult in active mice due to a high proportion of premature responses, a phenomenon much less pronounced in passive mice.

In previous delayed-response tasks involving unilateral texture presentation, we showed that short-term memory correlates diverged depending on behavioral strategy: passive mice recruited area P, whereas active mice recruited M2. Activity in area P reflected past sensory information, whereas M2 activity was more closely related to future action. Importantly, in the present DELAY task, information about the future action is unavailable during the delay because the second texture had not yet been presented. This explains why frontal areas such as M2 were not involved in maintaining or transferring WM in this task, and why area P emerged as a more suitable substrate for interhemispheric WM transfer.

When faced with a delay, mice that adopted an active strategy appeared to compensate by increasing movement and relying on BC-mediated transfer, despite the inability of BC to stably maintain WM. In contrast, a passive strategy promoted interhemispheric transfer via P and supported more effective WM maintenance. Thus, while an active strategy favors rapid sensory exchange through BC, it is poorly suited for delayed comparison, whereas a passive strategy supports stable interhemispheric WM transfer.

### Area P as a central node in maintaining and transferring higher-order sensory information

This study highlights the importance of area P, including surrounding ventral regions such as LM and POR, in transferring higher-order sensory information across hemispheres. This extends previous work implicating area P in short-term memory, learning, and higher-order sensory processing, particularly in the visual domain ^32–34,39,52–55^. We further found that area P was unilaterally active prior to stimulus presentation exclusively in passive mice (Fig. S4), suggesting a role in anticipation and preparation for interhemispheric comparison.

Area P is well positioned anatomically to serve this function. It lies along the ventral processing stream and is reciprocally connected with parahippocampal and entorhinal cortices, striatum, M2, and other association areas^38,56,57^, making it a major hub linking prior experience with current sensory input. Its strong callosal projections to contralateral P, together with its proximity to sensory cortices of multiple modalities, make it an ideal candidate for higher-order interhemispheric transfer.

The selective recruitment of area P in passive mice suggests that this region is highly plastic and engaged on demand, particularly when tasks require the integration and maintenance of higher-order information such as WM. When simpler sensory information must be transferred rapidly, other regions such as BC may instead mediate interhemispheric exchange^6,22,29,30,42,58^.

Area P may be analogous to multisensory association regions in humans and non-human primates, such as the lateral occipital cortex^59–61^. In contrast, frontal regions including anterior lateral motor (ALM) cortex and M2 have been strongly implicated in WM maintenance when future actions can be anticipated^31,62–66^. In the present task, the delay period was placed between sensory stimuli such that the upcoming motor choice was unknown, thereby isolating sensory-related WM. Interhemispheric connections in frontal cortex may therefore be engaged primarily during coordinated bilateral movements, action planning, or robust maintenance of motor choices that reverberate between both ALMs^67^. In summary, area P plays a central role in higher-order multisensory processing and interhemispheric transfer, enabling the integration of sensory information across time and hemispheres and supporting a unified percept of the external world.

## Methods

### Animals

A total of five adult male mice (8-36 weeks old) were used in this study. All experiments were approved by the Institutional Animal Care and Use Committee (IACUC) at the Hebrew University of Jerusalem, Israel (Permit Number: MD-20-16065-4). All mice expressed a calcium indicator (GCaMP6f) across the entire dorsal cortex. Two mice were triple-transgenic (Rasgrf2-2A-dCre;CamK2a-tTA;TITL-GCaMP6f;^32–34,68–71^) and three C57BL/6 mice injected with pAAV.Syn.GCaMP6f.WPRE.SV40 (AAV9 serotype; Addgene) into 20 cortical sites targeting the superficial layers of the neocortex^72^. Both methods resulted in homogeneous expression across the superficial layers of the dorsal cortex.

### Wide-field preparation for cortex-wide imaging

We used an intact skull preparation to enable chronic wide-field calcium imaging of neocortical activity over several months^32–34,53,70,71,73,74^. Mice were anesthetized with 2% isoflurane (in pure O₂), and body temperature was maintained at 37°C. Local analgesia (1% lidocaine) was applied, the skull was exposed and cleaned, and the overlying muscles were removed to allow access to the entire dorsal surface of both hemispheres (∼12 × 8 mm², extending from approximately 3 mm anterior to bregma to ∼1 mm posterior to lambda, and from the midline to at least 5 mm laterally). In C57BL/6 mice (n=3), following skull exposure, 200 nl of the AAV virus was injected superficially at 20 evenly spaced sites along the dorsal cortical surface of both hemispheres using a microinjector (World Precision Instruments). Stereotaxic coordinates, including injection angles and depths, were determined based on the Paxinos and Franklin Mouse Brain Atlas (4th Edition, 2013) to target cortical layer 2/3. All coordinates were referenced to bregma and the pial surface, as defined in the atlas. Injection sites were distributed to achieve broad bilateral coverage of the dorsal neocortex. Injection accuracy and targeting specificity were verified post hoc via histological analysis (Fig. 3a bottom). Next, for both transgenic and C57BL/6 mice, a wall was constructed around the imaging field using UV-curved adhesive (iBond) and dental cement “worms” (Charisma). Transparent dental cement (Tetric EvoFlow T1) was then applied homogeneously over the exposed skull. Finally, a metal post for head fixation was attached to the posterior part of the skull, centered between the hemispheres. This minimally invasive preparation enabled high-quality, daily imaging over extended periods, with a high success rate.

#### Behavioral tasks

Following recovery, mice underwent one week of handling and habituation to head fixation. Subsequently, they were trained for one day to lick in response to texture presentation. All mice were trained and imaged in two tasks, beginning with the BOTH task and subsequently progressing to the DELAY task.

#### BOTH task

To study interhemispheric transfer of sensory information, the animals were trained on a novel matching task (termed BOTH task) in which they were required to match between two textures presented simultaneously to both whisker pads. Each trial began with an auditory cue (two 2 kHz beeps, each 100 ms in duration with a 50 ms interval), signaling the simultaneous approach of two sandpaper textures (grit size P100: rough; P1200: smooth) to both sides of the mouse’s whiskers (Fig. 1a; Videos S1, 2). There were four possible stimulus configurations: P100 left and P100 right (P100P100); P1200 left and P1200 right (P1200P1200); P100 left and P1200 right (P100P1200); P1200 left and P100 right (P1200P100). Match trials with identical textures on both sides (i.e., P100P100 and P1200P1200), were defined a ‘go’ condition. Non-match trials differing in textures (i.e., P100P1200 and P1200P100), were defined as the no-go condition. Trial types were presented in a pseudo-randomized order with no more than three repetitions of the same condition. Textures remained in contact with the whiskers for 2 seconds. Following this period, the textures were retracted, and mice were rewarded with water for licking in go trials (i.e., Hit trial) with a 2 second response window. In no-go trials, licking responses (false alarms; FA) triggered mild white noise as punishment. No reward or punishment was given when mice correctly withheld licking in no-go trials (correct rejections; CR) or failed to lick in go trials (miss). Each trial was followed by a 4 second inter-trial interval.

This task design required mice to generalize matching textures, that is to lick for both go conditions and withhold licking for both no-go conditions. We observed mostly no significance performance bias between go or no-go performance (p>0.05; Signed rank test for each mouse; Hit rate P100P100 vs P1200P1200; CR rate between P100P1200 vs. P1200P100; Fig. S1). Consequently, successful performance requires identifying the texture-type on each side and comparing sensory information across hemispheres. Specifically, the design enables a dissociation of the texture-type from the choice of the mouse, for example by comparing between the two Hit trials.

The training protocol followed standard go/no-go paradigms^34,53,70,72,75^. Initially, both ‘go’ conditions were presented with equal probability. After 50 trials, the probability of both ‘no-go’ conditions was gradually increased by 10% every 50 trials, reaching 50% within the first two days of training (i.e., 0.25 probability for each stimulus pair). At that point, most mice continuously licked for all conditions regardless of the outcome. After around 100 trials, we increased no-go probability to 80% and waited for mice to perform three continuous CR trials before returning to a 50% probability. This was done for several rounds until mice increased their performance, specifically withheld licking for both no-go conditions. In mice that continued to lick for both textures, we additionally repeated the wrong trial until a correct response was made. In addition, in mice that developed a bias towards certain stimulus pairs, we counterbalanced the stimulus probability by decreasing the probability for the biased pairs. In all mice, a 50% protocol was presented with no repetitions, as soon as they reached an expert level (d′ > 1.5). Training duration for the BOTH task ranged from 11 to 31 training days. Upon reaching expert performance, wide-field imaging of the entire dorsal cortex (dual-hemispheres) was performed during task performance. We collected 11-16 imaging days per mouse. All training and imaging were conducted in complete darkness.

#### DELAY task

Following imaging in the BOTH task, all mice continued training on a DELAY task in which we introduced a delay period between the two textures (Fig. 1d; Videos S3, 4). The first stimulus was presented to the right whisker pad for 2 seconds after which it was retracted and a 2 seconds delay period was introduced. Subsequently, the second stimulus was presented on the left whisker pad for 2 seconds. Only after the second presentation was there a 2 seconds reward period. If the two stimuli matched, the trial was considered a go trial, and the mouse was required to lick to receive a reward (Hit). If the textures did not match, the trial was considered a no-go trial, and the mouse had to withhold licking (CR). This sequential presentation introduced a higher cognitive demand: the mouse had to identify the first texture-type, retain it in WM, presumably transfer that information to the other hemisphere and compare it to the second incoming texture.

In terms of training, we increased the delay period by 100 ms where licking within the delay period resulted in a white-noise punishment (i.e., early licks). Delay duration was increased only if mice reduced early licks while maintaining high performance. Training duration for the DELAY task ranged from 11 to 27 additional days, and the threshold for expert performance was defined as d’=1. Upon reaching expert level, we imaged the dorsal cortex during task performance, obtaining 12-22 imaging sessions per mouse. In some mice we also performed cortex-wide imaging of both the BOTH and DELAY tasks on the same day with expert performance in both tasks. In total, training and imaging of both tasks ranged from 161-196 days (gross) for each mouse, across a cohort of five mice.

#### Wide-field imaging

The wide-field imaging setup^34,53^ consisted of a sensitive CMOS camera (Hamamatsu Orca Flash 4.0 v3) mounted above a dual-objective optical system. Two Navitar objectives (top: D-5095, 50 mm f/0.95; bottom, inverted: D-2595, 35 mm f/0.95) were aligned through a dichroic filter cube (510 nm; AHF; Beamsplitter T510LPXRXT, Thorlabs). This configuration provided a field of view that encompassed the entire dorsal surface of both cortical hemispheres. Excitation light from a blue LED (Thorlabs M470L3) passed through an excitation filter (480/40 nm BrightLine HC), a diffuser, and a collimator, was reflected by the dichroic mirror, and then focused through the bottom objective to approximately 100 μm below the cortical surface. Emitted green fluorescence passed through both objectives and an emission filter (514/30 nm BrightLine HC) before reaching the camera. The total blue light power on the preparation remained below 5 mW (< 0.1 mW/mm²), a level at which no photobleaching was observed. Data were acquired at a temporal resolution of 20 Hz and a spatial resolution of 512 × 512 pixels. Activation maps were registered onto the Mouse Brain Atlas based on skull coordinates and functional patches (© 2004 Allen Institute for Brain Science; Available from: https://mouse.brain-map.org/).

#### Control for Non-Calcium-Dependent Signals

All imaging in this study was performed using single-wavelength excitation at 473 nm to measure calcium dynamics, following previous studies^32–34,71^. However, this approach may also capture non-calcium-dependent signals, including hemodynamic responses, which could potentially confound the interpretation of fluorescence changes. To control non-calcium-dependent signals, 405 nm excitation light was used instead of 473 nm light in a subset of the imaging sessions. In these control sessions, we did not observed the activity patterns reported in our main results, consistent with previous studies from our lab (Fig. S5;^32,34,71^). Furthermore, since the calcium signals in this protocol are robust and relatively fast, correction for hemodynamic contamination does not substantially affect the signal^71^.

#### Body movement tracking

We used a body camera to detect general movements of the mouse (The Imaging Source DMK 33UX273; 30 Hz frame rate; Figs. 2, 5). For each imaging day, we first outlined the forelimbs and back (one area of interest for each), which were reliable areas for detecting general movements, similar to previous studies^32,34,53,70,72^. Next, we calculated the body movement (defined as 1 minus the frame-to-frame correlation) within these areas as a function of time for each trial. Thresholding at 3 s.d. above baseline (across time frames before stimulus cue) resulted in a binary movement vector (either ‘moving’ or ‘quiet’ for each time frame) for each trial.

## Data analysis

Data analysis was performed using MATLAB software (MathWorks). Wide-field fluorescence images were downsampled to 256 × 256 pixels, and pixels outside the imaging area were discarded. This resulted in a spatial resolution of ∼40 μm/pixel, which was sufficient to determine cortical borders, despite further scattering of emitted light through the tissue and skull. For each pixel and each trial, the ΔF/F was calculated by dividing the raw signal by the baseline signal averaged over several frames before the stimulus cue (frame 0 division). Each trial was labeled and the correct body vector was assigned based on the texture stop position, which was clearly visible in the body camera in all trials.

### Behavior analysis

We defined the behavioral strategy of the mouse based on the movement vector for each trial that was derived from the body camera. In the BOTH task, we defined activeness as the average movement probability during the sensation period (−1 to +1 seconds relative to texture stop) in Hit trials (Fig. 2c). This was done for each mouse and each recording session separately. In addition, we calculated the movement onset for each recording session as the peak of the 2^nd^ derivative of the movement probability vector (Fig. 2d; movement probability vectors presented in Fig. 2b). Based on these two parameters, activeness and movement onset, we divided the five mice into passive (n = 3; low activeness and late movement onset) and active (n = 2; high activeness and early movement onset). These behavioral strategies are stable within each mouse but can also vary depending on other factors such as arousal and motivation. Previous studies have shown that behavioral strategies have a strong and long-lasting effect on cortex-wide dynamics during task performance^32–34^.

In the DELAY task, we defined an impatience index based on the movement vector for each mouse and recording session separately (Fig. 5d; movement probability vectors presented in Fig. 5b,c). The impatience index was defined as the movement probability during the first second of the delay period minus the movement probability one second before the start of the delay period (i.e., baseline). High impatience index values imply that the mouse started to move vigorously as the first texture moved out, not necessarily licking. Low impatience index values indicate that the mouse remained still as the first texture was retracted. Based on the impatience index, we were able to show that active mice (defined in the BOTH task) have high impatience index values whereas passive mice have low values.

### Correlation analysis

To study the relationship between different cortical areas with emphasis on inter-hemispheric relationships, we calculated the pairwise correlation (Pearson correlation coefficient; r) during the BOTH task (Figs. 4 and S6). We focused on interhemispheric correlations with the motivation to detect positive connections across hemispheres as the two textures simultaneously came into contact. Since single trials display rather large cortex-wide co-fluctuations, we calculated correlations between two pairs each from a different trial (i.e., trial-shuffled correlations). This procedure reduces within-trial co-fluctuations and extracts mainly dynamics that are time-locked across trials. This procedure is similar to calculating the signal correlation (i.e., correlation of the average signal) but maintains trial statistics. First, we calculated the seed correlation map focusing on three key areas: left P, left BC, and left M1. In this analysis, we correlated the responses in a chosen seed area (left P, left BC, or left M1) with each pixel in the imaged area (Fig. 4a, b). Each pixel in the map depicts the r values (trial shuffled and averaged across trials) between the seed area and that pixel. This can be done as a function of time with a running window (n = 20 frames, 1 second) to produce a seed correlation movie. Next, we calculated the full correlation matrix between all 20 areas in a similar fashion (i.e., trial-shuffled; running window) and focused on three interhemispheric homologous pairs: P-P, BC-BC, and M1-M1 (Fig. 4b-d). The full correlation matrix is presented in Figure S6e.

### Decoding analysis

To study how well cortex-wide dynamics decode the texture-type (P100 or P1200), we trained a support vector machine (SVM) to classify trials into the two different Hit conditions (i.e., Hit P100P100 vs. Hit P1200P1200) based on cortex-wide neuronal dynamics^32–34^. A linear SVM was trained on the neuronal dynamics of the 20 cortical areas during the BOTH and DELAY tasks (Figs. 5 and 8) for each time frame separately (80% of the trials; ridge regularization; cross-validation n = 10; equal size groups). The remaining 20% of the trials were used as a test set for the trained SVM model and accuracy was calculated. This was done for each mouse separately. A trial-shuffled distribution (n = 100 interation) was used as a control, and observed values exceeding 2 STD of the trial-shuffled distribution were considered significant (Fig. 5b). In addition, the weights assigned to each brain area by the classifier can also be plotted as a function of time (Fig. 5c). The full presentaion of accuracies and weights for each mouse is presented in Figures S7 and S9 for the BOTH and DELAY tasks, respectively.

### Switch analysis

During the DELAY task, we were interested in detecting the time of possible transfer of information from the left to the right hemisphere, which we termed as the time of switch. Given the results from the amplitude analysis, we focused on quantifying the time of switch between the left P and the right P in passive mice and between the left BC and the right BC in active mice, i.e., between two homologous areas from different hemispheres (Fig. 7e-g). Switch detection was calculated for each trial separately. Responses from both areas were first smoothed (Gaussian kernel with half-width = 21) and z-scored, and values below a z-score of 1 were discarded. The time of switch was defined as the first frame in which the right area had a higher response than the left area by z-score of 0.1 within a time window of -0.5 to 4 s relative to the stop of the first texture (i.e., including the first texture presentation and the delay period).

### Statistical analysis

In general, non-parametric two-tailed statistical tests were used: the Wilcoxon rank-sum test to compare two medians from two populations, or the Wilcoxon signed-rank test to compare a population’s median to zero (or between two paired populations). Multiple comparisons correction was used when comparing more than two groups. Trial-shuffled data were used to directly compute sample distributions. Importantly, we implemented an individual mouse approach where we statistically validate each mouse separately (based on its own recording sessions; n ≥ 11 for each mouse), displaying reproducibility across mice.

### Histology

To verify the location of GCaMP6f in 2/3 cortical layer of C57BL/6 mice, animals were deeply anesthetized with an overdose of Pental and transcardially perfused with phosphate-buffered saline (PBS), followed by 4% paraformaldehyde (PFA) in PBS. Brains were post-fixed for 12-24 hours in 4% PFA and subsequently cryoprotected for over 24 hours in 30% sucrose in PBS. Coronal sections (60-100 μm) of the entire brain were prepared using a vibratome (VT 1000S, Leica). Sections were mounted on glass slides, covered with mounting medium containing DAPI (SouthernBiotech) and coverslipped. Low-magnification images were acquired using a Nikon SMZ-25 fluorescent stereoscope equipped with 1× and 2× objectives (Fig. 3a). All 5 mice displayed a relatively homogenous expression of GCaMP6f in mostly superficial layers across the whole dorsal cortex in both hemispheres.

## Acknowledgements

We thank Malak Abumadi for assistance with genotyping and mouse colony management; Shaked Yadlin for support with SVM analysis; Tamar Licht for histology expertise; and Rotem Goldschmidt for help with the Allen Brain Atlas. We also thank Fritjof Helmchen and Philipp Bethge for helping with the transgenic mouse lines. This work is funded by the European Union (ERC Starting Grant, MESO-AG, 101040378) and a Hebrew University start-up grant.

## Author contributions

E.A. and A.G. conceptualized the study. E.A., S.D.S and A.G. performed the experiments and preprocessed the data. E.A. and A.G. analyzed the data. E.A. and A.G. wrote the manuscript. All authors read and approved the final version of the manuscript.

## Data availability

Data will be shared upon reasonable request by the corresponding author. No new material was generated in this work.

## Code availability

Custom codes for data analysis were written in MATLAB and are available from the corresponding author upon request.

## References

1. Robertson, L. C. Binding, spatial attention and perceptual awareness. Nat. Rev. Neurosci. 4, 93 (2003).

2. Brincat, S. L. & Miller, E. K. Cognitive independence and interactions between cerebral hemispheres. Neuropsychologia 212, 109153 (2025).

3. Ocklenburg, S. & Guo, Z. V. Cross-hemispheric communication: Insights on lateralized brain functions. Neuron 112, 1222–1234 (2024).

4. Mundorf, A. & Ocklenburg, S. Looking at Both Sides: Integrating Data From Both Hemispheres Is Crucial in Rodent Neuroscience. Eur. J. Neurosci. 62, (2025).

5. Gazzaniga, M. S. Cerebral specialization and interhemispheric communication. Does the corpus callosum enable the human condition? Brain 123, 1293–1326 (2000).

6. Brincat, S. L. et al. Interhemispheric transfer of working memories. Neuron 109, 1055–1066.e4 (2021).

7. Christophel, T. B., Klink, P. C., Spitzer, B., Roelfsema, P. R. & Haynes, J. D. The Distributed Nature of Working Memory. Trends Cogn. Sci. 21, 111–124 (2017).

8. Baddeley, A. Working Memory: Theories, Models, and Controversies - annurev-psych-120710-100422. Annu. Rev. Psychol. 63, 1–29 (2012).

9. Hoy, J. L., Yavorska, I., Wehr, M. & Niell, C. M. Vision Drives Accurate Approach Behavior during Prey Capture in Laboratory Mice. Curr. Biol. 26, 3046–3052 (2016).

10. Sato, T. R. et al. Interhemispherically dynamic representation of an eye movement-related activity in mouse frontal cortex. Elife 8, 1–28 (2019).

11. Rivera-Olvera, A. et al. The universe is asymmetric, the mouse brain too. Mol. Psychiatry 30, 489–496 (2025).

12. Innocenti, G. M. General Organization of Callosal Connections in the Cerebral Cortex. 291–353 (1986) doi:10.1007/978-1-4613-2149-1_9.

13. Jarbo, K., Verstynen, T. & Schneider, W. In vivo quantification of global connectivity in the human corpus callosum. Neuroimage 59, 1988–1996 (2012).

14. van der Knaap, L. J. & van der Ham, I. J. M. How does the corpus callosum mediate interhemispheric transfer? A review. Behav. Brain Res. 223, 211–221 (2011).

15. Bloom, J. S. & Hynd, G. W. The role of the corpus callosum in interhemispheric transfer of information: excitation or inhibition? Neuropsychol. Rev. 15, 59–71 (2005).

16. Moon, H. S., Vo, T. T., Im, G. H., Hong, S. J. & Kim, S. G. Interhemispheric resting-state functional connectivity correlates with spontaneous neural interactions. Proc. Natl. Acad. Sci. U. S. A. 122, e2505294122 (2025).

17. Roland, J. L. et al. On the role of the corpus callosum in interhemispheric functional connectivity in humans. Proc. Natl. Acad. Sci. U. S. A. 114, 13278–13283 (2017).

18. Deligianni, F., Centeno, M., Carmichael, D. W. & Clayden, J. D. Relating resting-state fMRI and EEG whole-brain connectomes across frequency bands. Front. Neurosci. 8, 98767 (2014).

19. Shimaoka, D., Steinmetz, N. A., Harris, K. D. & Carandini, M. The impact of bilateral ongoing activity on evoked responses in mouse cortex. Elife 8, (2019).

20. Cybulska-Klosowicz, A. & Kossut, M. Early-phase of learning enhances communication between brain hemispheres. Eur. J. Neurosci. 24, 1470–1476 (2006).

21. Debowska, W., Liguz-Lecznar, M. & Kossut, M. Bilateral plasticity of Vibrissae SII representation induced by classical conditioning in mice. J. Neurosci. 31, 5447–5453 (2011).

22. Oran, Y., Katz, Y., Sokoletsky, M., Malina, K. C. K. & Lampl, I. Reduction of corpus callosum activity during whisking leads to interhemispheric decorrelation. Nat. Commun. 12, (2021).

23. Li, L., Rema, V. & Ebner, F. F. Chronic suppression of activity in barrel field cortex downregulates sensory responses in contralateral barrel field cortex. J. Neurophysiol. 94, 3342–3356 (2005).

24. Zhang, Z. & Zagha, E. Motor cortex gates distractor stimulus encoding in sensory cortex. Nat. Commun. 2023 141 14, 2097- (2023).

25. Van Ede, F., De Lange, F. P. & Maris, E. Anticipation increases tactile stimulus processing in the ipsilateral primary somatosensory cortex. Cereb. Cortex 24, 2562–2571 (2014).

26. DeCosta-Fortune, T. M. et al. Repetitive microstimulation in rat primary somatosensory cortex (SI) strengthens the connection between homotopic sites in the opposite SI and leads to expression of previously ineffective input from the ipsilateral forelimb. Brain Res. 1732, (2020).

27. Petrus, E. et al. Interhemispheric plasticity is mediated by maximal potentiation of callosal inputs. Proc. Natl. Acad. Sci. U. S. A. 116, 6391–6396 (2019).

28. Calford, M. B. & Tweedale, R. Interhemispheric transfer of plasticity in the cerebral cortex. Science 249, 805–807 (1990).

29. Park, H., Keri, H. V. S., Yoo, C., Bi, C. & Pluta, S. R. Bilateral integration in somatosensory cortex is controlled by behavioral relevance. Nat. Neurosci. 28, 1300–1310 (2025).

30. Shuler, M. G., Krupa, D. J. & Nicolelis, M. A. L. Bilateral integration of whisker information in the primary somatosensory cortex of rats. J. Neurosci. 21, 5251–5261 (2001).

31. Yin, X., Wang, Y., Li, J. & Guo, Z. V. Lateralization of short-term memory in the frontal cortex. Cell Rep. 40, 111190 (2022).

32. Gilad, A., Gallero-Salas, Y., Groos, D. & Helmchen, F. Behavioral Strategy Determines Frontal or Posterior Location of Short-Term Memory in Neocortex. Neuron 0, (2018).

33. Gallero-Salas, Y. et al. Sensory and Behavioral Components of Neocortical Signal Flow in Discrimination Tasks with Short-Term Memory. Neuron 109, 135–148.e6 (2021).

34. Rokach, R. O., Marmor, O., Levy, Y. & Gilad, A. ’What’ and ‘where’ brain-wide pathways are dominated by internal strategies. bioRxiv 2025.03.23.644791 (2025) doi:10.1101/2025.03.23.644791.

35. Romo, R. & de Lafuente, V. Conversion of sensory signals into perceptual decisions. Prog. Neurobiol. 103, 41–75 (2013).

36. Liu, D. et al. Medial prefrontal activity during delay period contributes to learning of a working memory task. Science (80-.). (2014) doi:10.1126/science.1256573.

37. Daniel, T. A., Katz, J. S. & Robinson, J. L. Delayed match-to-sample in working memory: A BrainMap meta-analysis. Biol. Psychol. 120, 10 (2016).

38. Oh, S. W. et al. A mesoscale connectome of the mouse brain. Nature 508, 207–214 (2014).

39. Wang, Q., Gao, E. & Burkhalter, A. Gateways of ventral and dorsal streams in mouse visual cortex. J. Neurosci. 31, 1905–18 (2011).

40. O’Reilly, J. X. et al. Causal effect of disconnection lesions on interhemispheric functional connectivity in rhesus monkeys. Proc. Natl. Acad. Sci. U. S. A. 110, 13982–13987 (2013).

41. Sperry, R. W. Hemisphere deconnection and unity in conscious awareness. Am. Psychol. 23, 723–733 (1968).

42. Broschard, M. B., Roy, J. E., Brincat, S. L., Mahnke, M. K. & Miller, E. K. Evidence for an Active Handoff between Hemispheres during Target Tracking. J. Neurosci. 45, 1–11 (2025).

43. de León Reyes, N. S., Bragg-Gonzalo, L. & Nieto, M. Development and plasticity of the corpus callosum. Dev. 147, (2020).

44. Adaikkan, C. et al. Alterations in a cross-hemispheric circuit associates with novelty discrimination deficits in mouse models of neurodegeneration. Neuron 110, 3091–3105.e9 (2022).

45. Guo, Z. V. et al. Flow of cortical activity underlying a tactile decision in mice. Neuron 81, 179–94 (2014).

46. Pinto, L. et al. Task-Dependent Changes in the Large-Scale Dynamics and Necessity of Cortical Regions. Neuron 104, 810–824.e9 (2019).

47. Driscoll, L. N., Pettit, N. L., Minderer, M., Chettih, S. N. & Harvey, C. D. Dynamic Reorganization of Neuronal Activity Patterns in Parietal Cortex. Cell 170, 986–999.e16 (2017).

48. Steinmetz, N. A., Zatka-Haas, P., Carandini, M. & Harris, K. D. Distributed coding of choice, action and engagement across the mouse brain. Nature 576, 266–273 (2019).

49. Khilkevich, A. et al. Brain-wide dynamics linking sensation to action during decision-making. Nature 48–50 (2024) doi:10.1038/s41586-024-07908-w.

50. Chen, S. et al. Brain-wide neural activity underlying memory-guided movement. Cell 187, 676–691.e16 (2024).

51. Zhang, Y. et al. A brain-wide map of neural activity during complex behaviour. Nature 645, 177–191 (2025).

52. Consorti, A. et al. An essential role for the latero-medial secondary visual cortex in the acquisition and retention of visual perceptual learning in mice. Nat. Commun. 15, 1–16 (2024).

53. Pollak, Y. E., Sachdev, R., Larkum, M. & Gilad, A. Cortex-wide laminar dynamics diverge during learning. bioRxiv 2025.07.15.664840 (2025) doi:10.1101/2025.07.15.664840.

54. Glickfeld, L. L. & Olsen, S. R. Higher-Order Areas of the Mouse Visual Cortex. Annu. Rev. Vis. Sci. 3, 251–273 (2017).

55. Goltstein, P. M., Reinert, S., Bonhoeffer, T. & Hübener, M. Mouse visual cortex areas represent perceptual and semantic features of learned visual categories. Nat. Neurosci. 24, 1441–1451 (2021).

56. Wang, Q. & Burkhalter, A. Area map of mouse visual cortex. J. Comp. Neurol. 502, 339–357 (2007).

57. Wang, Q., Sporns, O. & Burkhalter, A. Network Analysis of Corticocortical Connections Reveals Ventral and Dorsal Processing Streams in Mouse Visual Cortex. J. Neurosci. 32, 4386–4399 (2012).

58. Cavanagh, P. & Alvarez, G. A. Tracking multiple targets with multifocal attention. Trends Cogn. Sci. 9, 349–354 (2005).

59. Amedi, A., Jacobson, G., Hendler, T., Malach, R. & Zohary, E. Convergence of Visual and Tactile Shape Processing in the Human Lateral Occipital Complex. 1202–1212 (2002).

60. Grill-Spector, K., Kourtzi, Z. & Kanwisher, N. The lateral occipital complex and its role in object recognition. Vision Res. 41, 1409–1422 (2001).

61. Orban, G. A., Van Essen, D. & Vanduffel, W. Comparative mapping of higher visual areas in monkeys and humans. Trends Cogn. Sci. 8, 315–324 (2004).

62. Li, N., Chen, T.-W., Guo, Z. V., Gerfen, C. R. & Svoboda, K. A motor cortex circuit for motor planning and movement. Nature 519, 51–56 (2015).

63. Chen, T.-W., Li, N., Daie, K. & Svoboda, K. A Map of Anticipatory Activity in Mouse Motor Cortex. Neuron 94, 866–879.e4 (2017).

64. Makino, H. et al. Transformation of Cortex-wide Emergent Properties during Motor Learning. Neuron 94, 880–890.e8 (2017).

65. Esmaeili, V. et al. Rapid suppression and sustained activation of distinct cortical regions for a delayed sensory-triggered motor response. Neuron 109, 2183–2201.e9 (2021).

66. Chen, G., Kang, B., Lindsey, J., Druckmann, S. & Li, N. Modularity and robustness of frontal cortical networks. Cell 184, 3717–3730.e24 (2021).

67. Li, N., Daie, K., Svoboda, K. & Druckmann, S. Robust neuronal dynamics in premotor cortex during motor planning. Nature 532, 459–64 (2016).

68. Harris, J. A. et al. Anatomical characterization of Cre driver mice for neural circuit mapping and manipulation. Front. Neural Circuits 8, 1–16 (2014).

69. Madisen, L. et al. Transgenic mice for intersectional targeting of neural sensors and effectors with high specificity and performance. Neuron 85, 942–958 (2015).

70. Gilad, A. & Helmchen, F. Spatiotemporal refinement of signal flow through association cortex during learning. Nat. Commun. 11, 1–14 (2020).

71. Marmor, O., Pollak, Y., Doron, C., Helmchen, F. & Gilad, A. History information emerges in the cortex during learning. Elife 12, (2023).

72. Levitan, D. & Gilad, A. Amygdala and Cortex Relationships during Learning of a Sensory Discrimination Task. J. Neurosci. 44, 1–13 (2024).

73. Silasi, G., Xiao, D., Vanni, M. P., Chen, A. C. N. & Murphy, T. H. Intact skull chronic windows for mesoscopic wide-field imaging in awake mice. J. Neurosci. Methods 267, 141–149 (2016).

74. Gilad, A. Wide-field imaging in behaving mice as a tool to study cognitive function. Neurophotonics 11, 1–16 (2024).

75. Chen, J. L., Carta, S., Soldado-Magraner, J., Schneider, B. L. & Helmchen, F. Behaviour-dependent recruitment of long-range projection neurons in somatosensory cortex. Nature 499, 336–40 (2013).

