## Supplementary figures for "Interhemispheric transfer of sensory and working memory information is dictated by behavioral strategy"

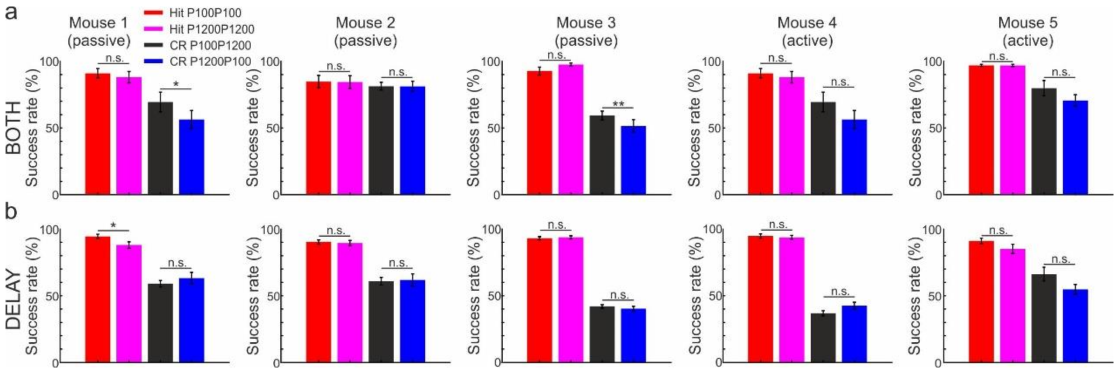


**Figure S1. Behavioral performance in each condition during the BOTH and DELAY tasks. a.** Success rate for each condition (i.e., two Hit conditions and two CR conditions) during the BOTH task for each mouse. Error bars depicts SEM across recording sessions (n=13, 16, 13, 12 and 11 for mouse 1-5 respectively). **b.** Same as a but for the DELAY task (n=24, 44, 32, 24 and 26 for mouse 1-5 respectively).

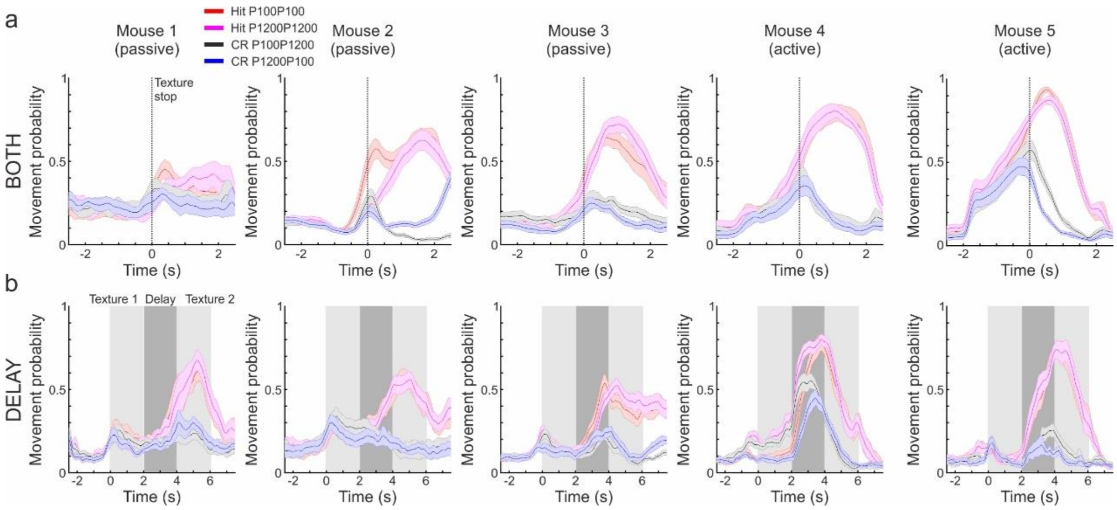


**Figure S2. Movement probability for each condition during the tasks. a.** Movement probability for each condition (i.e., two Hit conditions and two CR conditions) and each mouse during the BOTH task. Error bars depicts SEM across recording sessions (n=13, 16, 13, 12 and 11 for mouse 1-5 respectively). **b.** Same as a but for the DELAY task (n=24, 44, 32, 24 and 26 for mouse 1-5 respectively).

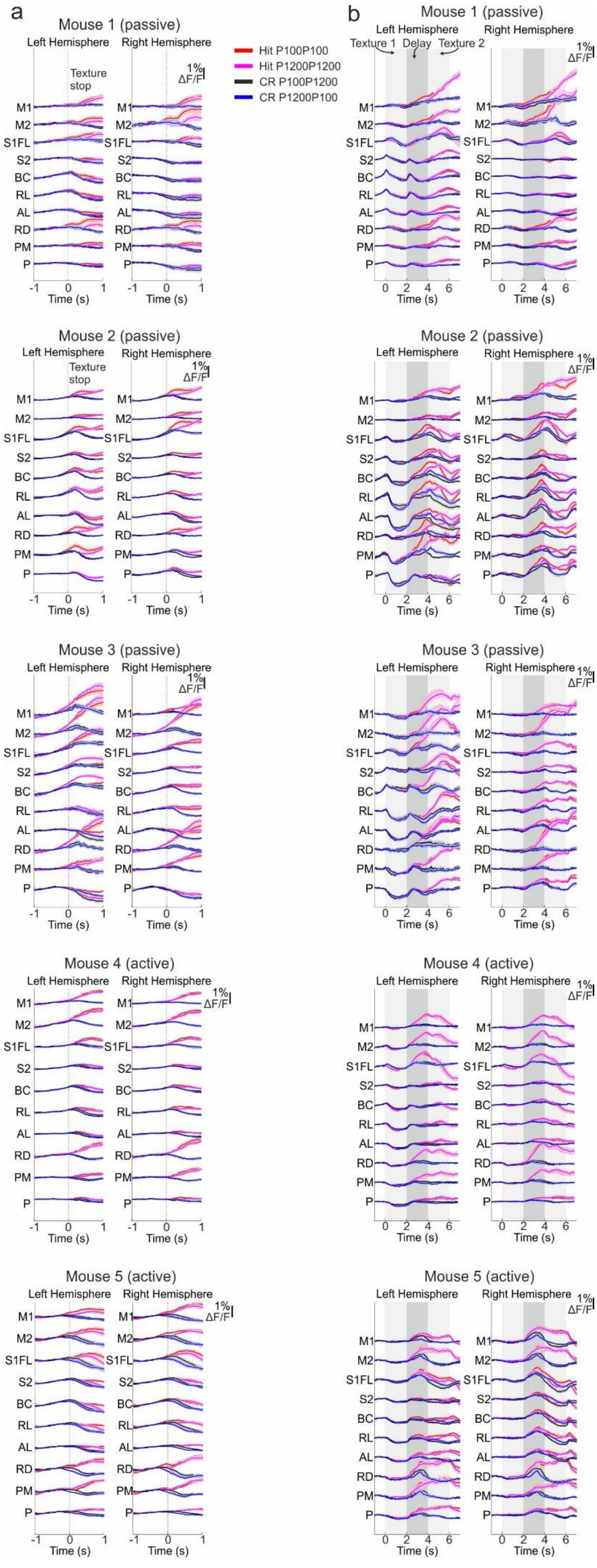
a
Mouse 1 (passive)
Mouse 2 (passive)
Mouse 3 (passive)
Mouse 4 (active)
Mouse 5 (active)
Left Hemisphere
Right Hemisphere
Texture stop
1% ΔF/F
Hit P100P100
Hit P1200P1200
CR P100P1200
CR P1200P100
M1
M2
S1FL
S2
BC
RL
AL
RD
PM
P
Time (s)
b
Texture 1
Delay
Texture 2

**Figure S3 Response profiles during the BOTH and DELAY tasks. a.** Temporal response profiles during the BOTH task for each cortical area (n=20) and each condition (2 Hit conditions and 2 CR conditions) plotted for each mouse separately. Error bars depict SEM across recording sessions (n=13, 16, 13, 12 and 11 for mouse 1-5 respectively). **b.** Same as in a, but for the DELAY task. Error bar depict SEM across recording sessions (n=24, 44, 32, 24 and 26 for mouse 1-5 respectively).

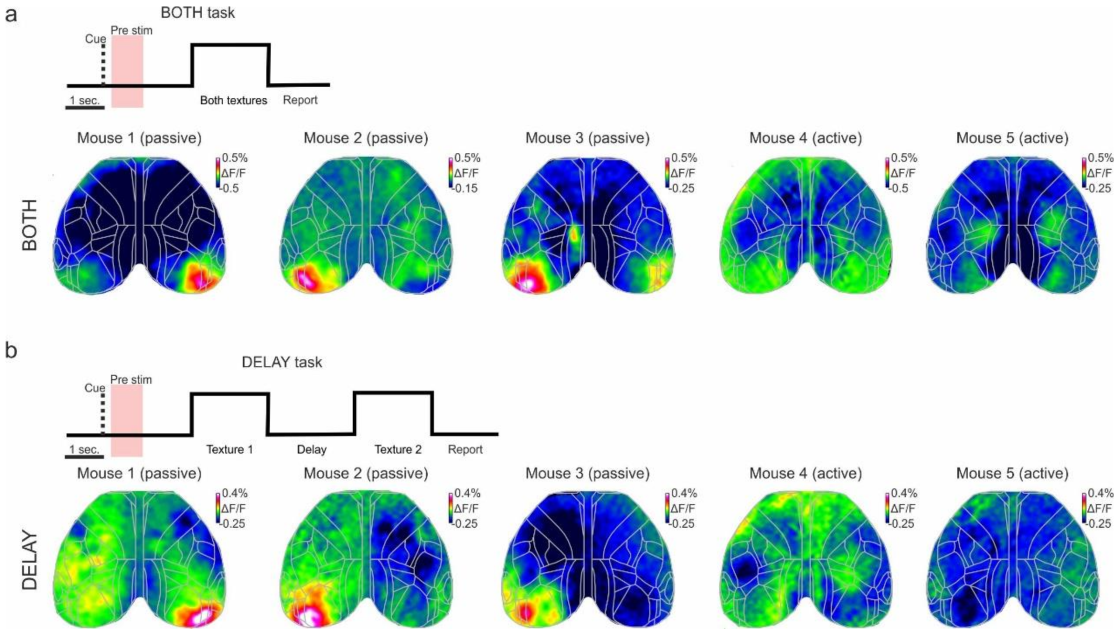


**Figure S4. Area P is activated before the stimulus only in passive mice. a.** Example activity maps (ΔF/F) before the sensation period (Pre-stim; -1.5 to -1 before texture stop; Pink shaded area) for each mouse. Color denotes ΔF/F. Only passive mice display a prominent activity patch in one of the P areas. **b.** Same as in b, but during the DELAY task.

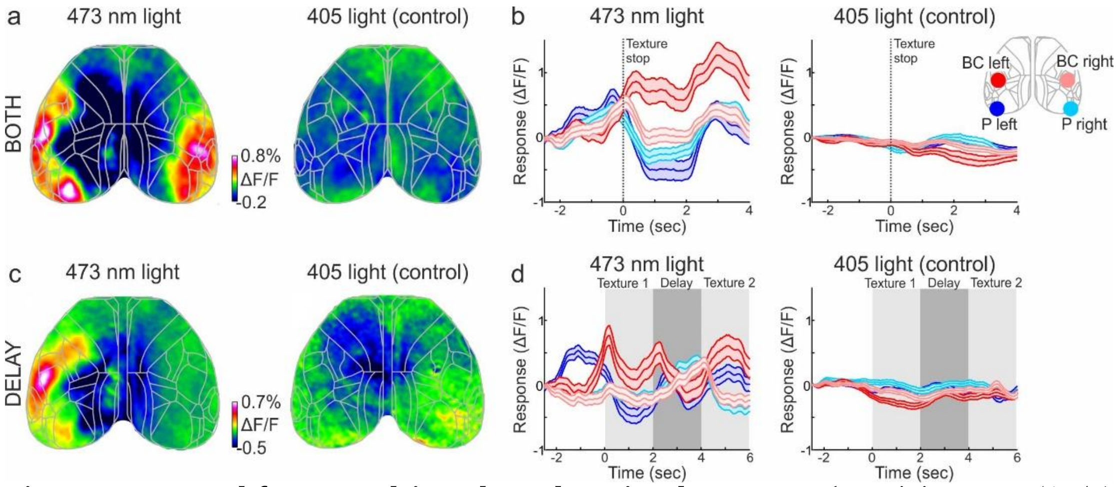


**Figure S5. Control for non-calcium dependent signals. a.** Example activity maps (ΔF/F) during the sensation period in the BOTH task for the regular 473 nm light (left) and a control 405 nm (right; Same passive mouse). **b.** Temporal responses for the example in a in four cortical areas (left and right P; Left and right BC) for the 473 (left) and 405 (right) lights. Error bars depict SEM across trials (n=45 and 35 trials for the 473 and 405 light respectively) **c, d.** Same as a and b but during the DELAY task (n=68 and 44 trials for the 473 and 405 light respectively).

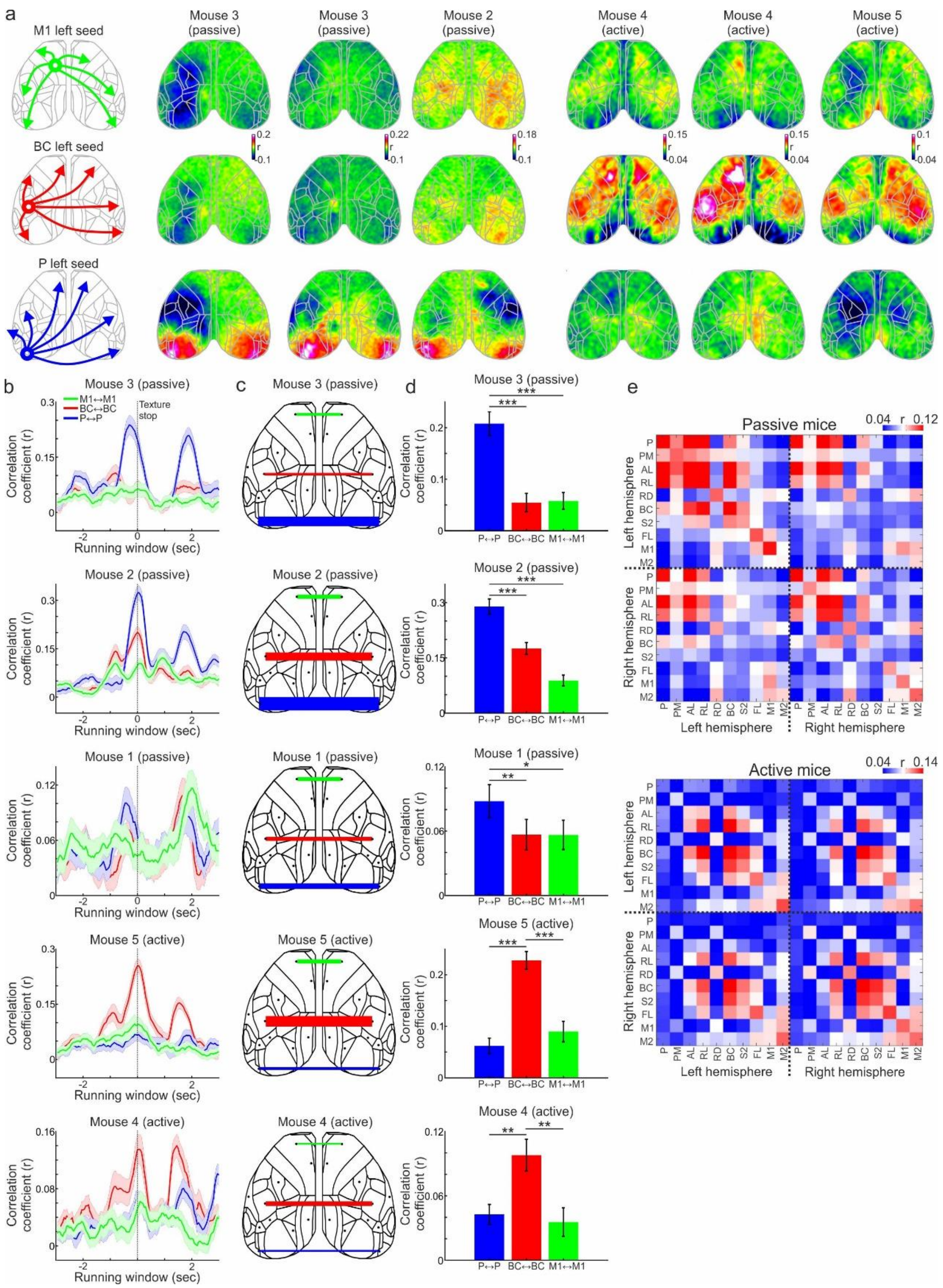


**Figure S6. Correlations during the BOTH task. a.** *Left:* schematic illustration for three different seed correlation maps (for a window during the sensation period) each displaying a different seed area (M1, BC and P). *Middle:* Corresponding seed correlation maps (colors depict r values) in an example passive mouse. *Right:* Example active mouse. **b.** Correlations (; see Methods) in a running window (1 sec.) for three interhemispheric pairs (P and P in blue; BC and BC in red; M1 and M1 in green) for each passive and active mouse. Error bars depict SEM across recording sessions n=13, 16, 13, 12 and 11 for mouse 1-5 respectively). **c.** Correlation values (r) of the three interhemispheric pairs during the sensation period plotted on top of the cortical map for an example passive (top) and passive (bottom) mouse. Thick lines depict high r values. **d.** Correlation values for the three interhemispheric pairs during the sensation period for each mouse. Error bars as in b. **e.** Full correlation matrices between all pairs in passive (top) and active (bottom) mice. color depict r values. * - $p<0.05$. ** - $p<0.01$. *** - $p<0.001$. Wilcoxon rank sum test.

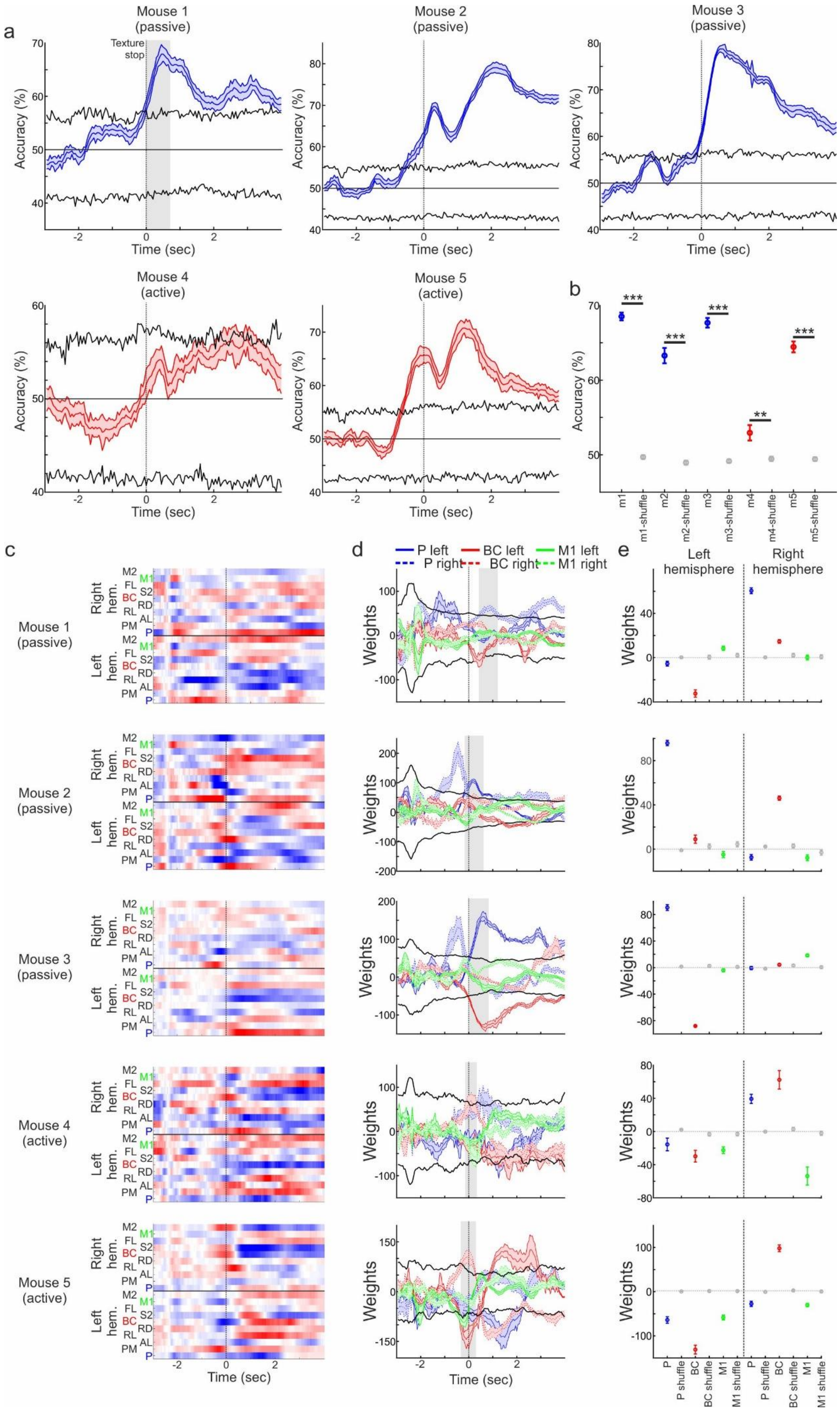

a
Mouse 1 (passive)
Texture stop
Mouse 2 (passive)
Mouse 3 (passive)
Mouse 4 (active)
Mouse 5 (active)
Accuracy (%)
Time (sec)
b
m1
m1-shuffle
m2
m2-shuffle
m3
m3-shuffle
m4
m4-shuffle
m5
m5-shuffle
c
Mouse 1 (passive)
Mouse 2 (passive)
Mouse 3 (passive)
Mouse 4 (active)
Mouse 5 (active)
Right hem.
Left hem.
M2
M1
FL
S2
BC
RD
RL
AL
PM
P
d
P left
BC left
M1 left
P right
BC right
M1 right
Weights
e
Left hemisphere
Right hemisphere
P
P shuffle
BC
BC shuffle
M1
M1 shuffle

**Figure S7. SVM products during the BOTH task for each mouse. a.** Accuracy as a function of time of an SVM classifier for texture types (i.e., between the two Hit trials: P100P100 vs. P1200P1200) during the BOTH task for each passive (blue) and active (red) mouse. Error bars depict SEM of cross validations (n=10). Black lines depict mean±2*STD of trial shuffled distribution. **b.** Mean accuracy during the sensation period for each mouse. In gray are trial shuffled distributions (n=100). **c.** Temporal profiles of the weights assigned to all cortical areas in each mouse. Red and blue indicate positive and negative weights respectively. **d.** Temporal profiles of weights the classifier assigned to areas P (blue), BC (red) and M1 (green). Solid lines are left hemisphere and dashed lines are left hemisphere. Error bars as in b. **e.** Corresponding weights for each mouse and the 6 different cortical areas along with their trial shuffled weights in gray. * - $p<0.05$. ** - $p<0.01$. *** - $p<0.001$. Wilcoxon rank sum test.

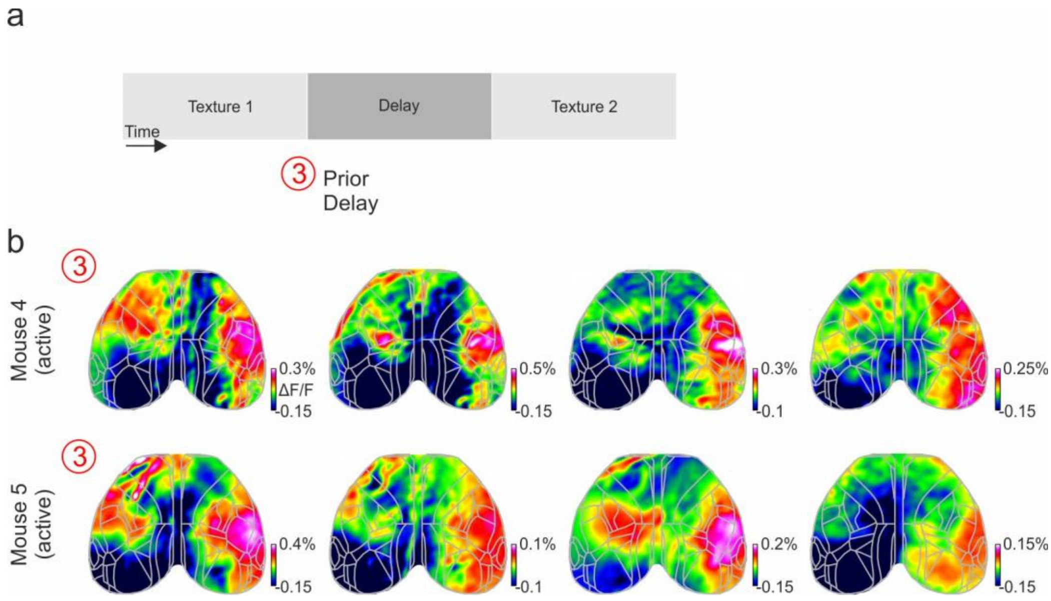


**Figure S8. Example activity maps in active mice during the DELAY task. a.** Definition of the prior delay periods during the DELAY task in active mice. **b.** Example activity maps (ΔF/F) from different recording days during the early delay period (1.5 to 2 seconds after the first texture stop) for the two active mice (mice 4 and 5).

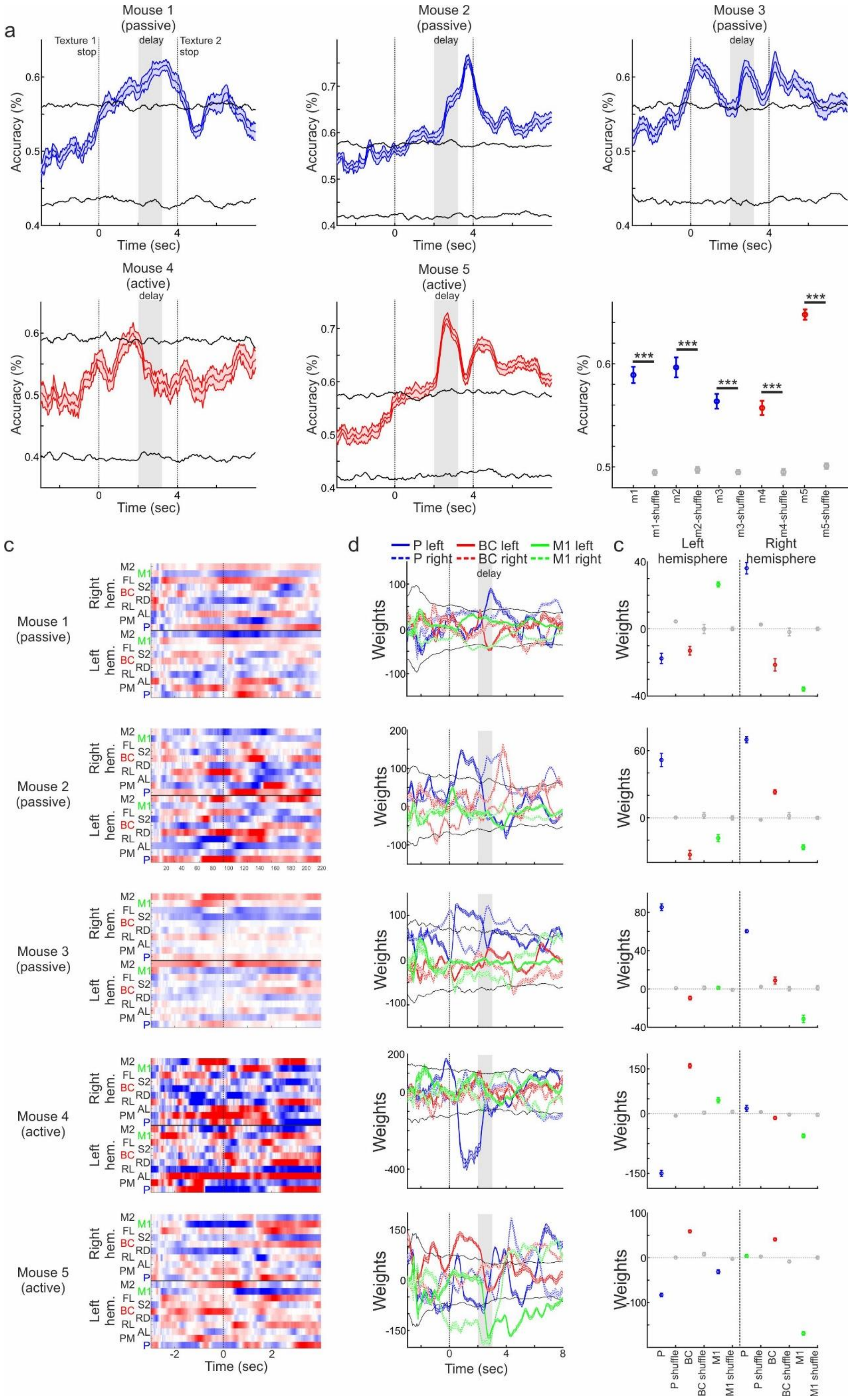

a
Mouse 1 (passive)
Mouse 2 (passive)
Mouse 3 (passive)
Mouse 4 (active)
Mouse 5 (active)
Texture 1 stop
delay
Texture 2 stop
Accuracy (%)
Time (sec)
***
m1
m1-shuffle
m2
m2-shuffle
m3
m3-shuffle
m4
m4-shuffle
m5
m5-shuffle
c
d
P left
P right
BC left
BC right
M1 left
M1 right
Left hemisphere
Right hemisphere
Right hem.
Left hem.
M2
M1
FL
S2
BC
RD
RL
AL
PM
P
Weights
P shuffle
BC shuffle
M1 shuffle

**Figure S9. SVM products during the DELAY task for each mouse. a.** Accuracy as a function of time of an SVM classifier for texture types (i.e., between the two Hit trials: P100P100 vs. P1200P1200) during the DELAY task for each passive (blue) and active (red) mouse. Error bars depict SEM of cross validations (n=10). Black lines depict mean±2*STD of trial shuffled distribution. **b.** Mean accuracy during the sensation period for each mouse. In gray are trial shuffled distributions (n=100). **c.** Temporal profiles of the weights assigned to all cortical areas in each mouse. Red and blue indicate positive and negative weights respectively. **d.** Temporal profiles of weights the classifier assigned to areas P (blue), BC (red) and M1 (green). Solid lines are left hemisphere and dashed lines are right hemisphere. Error bars as in *b*. **e.** Corresponding weights for each mouse and the 6 different cortical areas along with their trial shuffled weights in gray. * - $p<0.05$. ** - $p<0.01$. *** - $p<0.001$. Wilcoxon rank sum test.

**Video S1.** Video of body camera in several trials for a passive mouse during the BOTH task

**Video S2.** Video of body camera in several trials for an active mouse during the BOTH task

**Video S3.** Video of body camera in several trials for a passive mouse during the DELAY task

**Video S4.** Video of body camera in several trials for an active mouse during the DELAY task

**Video S5.** Activity map as a function of time in an example recording session for a passive mouse during the DELAY task (CRP100P1200). Color denotes $\Delta F/F$. Time 0 is the stop of the first texture. Notice the flow from left P to right P during the delay periods. Also notice the flows from BC to P in the left hemisphere just before the delay and the floe from P to BC in the right hemisphere toward the end of the delay period.

**Video S6.** Activity map as a function of time in an example recording session for an active mouse during the DELAY task (CRP100P1200). Color denotes $\Delta F/F$. Time 0 is the stop of the first texture. Notice the early flow across hemispheres from BC to BC within the first texture presentation.
